# The flagellar substrate specificity switch protein FlhB assembles onto the extra-membrane export gate to regulate type three secretion

**DOI:** 10.1101/686782

**Authors:** Lucas Kuhlen, Steven Johnson, Andreas Zeitler, Sandra Bäurle, Justin C. Deme, Rebecca Debo, Joseph Fisher, Samuel Wagner, Susan M. Lea

**Affiliations:** Sir William Dunn School of Pathology, University of Oxford, Oxford OX13RE, UK; Department of Chemistry, University of Oxford, Oxford, UK; University of Tübingen, Interfaculty Institute of Microbiology and Infection Medicine (IMIT), Elfriede-Aulhorn-Str. 6, 72076 Tübingen, Germany; Central Oxford Structural Microscopy and Imaging Centre, University of Oxford, Oxford OX1 3RE, UK; German Center for Infection Research (DZIF), partner-site Tübingen, Elfriede-Aulhorn-Str. 6, 72076 Tübingen, Germany; University Hospital Tübingen, Department of Thoracic & Cardiovascular Surgery, 72076 Tübingen, Germany; University of Tübingen Department of Geosciences, Centre for Applied Geosciences (ZAG), 72076 Tübingen, Germany

## Abstract

Export of proteins through type three secretion systems (T3SS) is critical for motility and virulence of many major bacterial pathogens. Proteins are transported through an export gate complex consisting of three proteins (FliPQR in flagellar systems, SctRST in virulence systems) that were initially annotated as membrane proteins, but which we have recently shown assemble into an extra-membranous helical assembly. A fourth putative membrane protein (FlhB/SctU) is essential to the export process, and also functions to “switch” secretion substrate specificity once the growing hook/needle structures reach their determined length. Here we present the structure of an export gate containing the switch protein from a *Vibrio* polar flagellar system at 3.2 Å resolution by cryo-electron microscopy. The structure reveals that the FlhB/SctU further extends the helical export gate assembly with its four putative transmembrane helices adopting an out-of-membrane location, wrapped around the other export gate components at the base of the structure. The unusual topology of the switch protein helices creates a loop that wraps around the bottom of the closed export gate complex. Structure-informed mutagenesis suggests that this loop is critical in gating secretion and we propose that a series of conformational changes in the type 3 secretion system trigger opening of the export gate through the interactions between FlhB/SctU and FliPQR/SctRST.

## Main

Type three secretion is a mechanism of bacterial protein secretion across both inner and outer bacterial membranes. It is found in the virulence-associated injectisome (vT3SS), a molecular syringe, and the bacterial flagellum (fT3SS), a motility organelle (*1*). Both families contribute in significant ways to bacterial pathogenesis. vT3SS facilitate secretion not only across the bacterial envelope but also insert translocon proteins at the tip of the needle into the eukaryotic host plasma membrane, allowing direct injection of virulence factors in the host cytoplasm. The fT3SS is responsible for construction of the flagellar filament in both Gram-negative and Gram-positive bacteria, and hence imparts pathogenicity (*2*) for example via the ability to swim towards favourable environments or sense environmental conditions (*3*).

T3SS are multi-megadalton protein complexes that are capable of bridging from the bacterial cytoplasm to the extracellular space. At the core of the secretion system is the highly conserved export apparatus (EA) (*4, 5*), which is made up of five predicted transmembrane (TM) proteins (SctR, SctS, SctT, SctU and SctV in the vT3SS; FliP, FliQ, FliR, FlhB and FlhA in the fT3SS). FlhA/SctV has been shown to form a nonameric ring (*6*), consisting of a large cytoplasmic domain situated below a hydrophobic domain predicted to contain 72 helices. This structure was proposed to surround an “export gate” through which substrates would enter the secretion pathway. This export gate is constructed from the other 4 EA proteins and was predicted to lie in the inner membrane. However, our recently determined structures of the *S. enterica* serovar Typhimurium FliP_5_Q_4_R_1_ and the *Shigella flexneri* SctR_5_S_4_T_1_ complexes (*7, 8*) demonstrated that the export gate is actually embedded within the proteinaceous core of the T3SS basal body, placing it above the predicted location of the inner membrane. Furthermore, the helical structure of the export gate makes it likely that it is responsible for nucleating the helical filaments that assemble above it (*9*). Interestingly, the export gate complex has also recently been proposed to facilitate inward transport across the inner membrane associated with nanotubes (*10, 11*). The final component of the EA, FlhB/SctU, has long been known to be essential for all T3SS-mediated protein secretion. In addition, FlhB/SctU has a major role in switching the specificity of secretion substrates, mediating the transition from the early components necessary to build the flagellar hook in fT3SS and injectisome needle in vT3SS, to the later subunits required for flagellar filament or injectisome translocon assembly. The FlhB/SctU family of proteins all contain an N-terminal hydrophobic sequence that is predicted to form 4 TM helices (FlhB_TM_) and a smaller cytoplasmic C-terminal domain (FlhB_C_). Crystal structures of the FlhB/SctU cytoplasmic domain from a range of species and systems (*12-14*) demonstrated a compact fold with an unusual autocatalytic cleavage site in a conserved NPTH sequence. Cleavage between the Asn and Pro residues, splitting FlhB_C_ into FlhB_CN_ and FlhB_CC_, is required for the switching event to occur and a variety of mechanisms have been proposed to explain the need for this unusual mechanism (*15*).

Little was known about the predicted TM portion of FlhB/SctU. Co-evolution analysis and molecular modelling led to suggestions that it forms a 4-helix bundle in the membrane (*16*), while crosslinks (*17*) and partial co-purification of FlhB with FliPQR were consistent with FlhB/SctU interacting with the export gate via a conserved site on the FliP_5_Q_4_R_1_ complex (*7*). However, given the inaccuracy of the TM predictions for the other export gate components revealed by the PQR structure, we sought to determine the molecular basis of the interaction of FlhB with FliPQR. Here we present the structure of the TM region of FlhB bound to the FliPQR complex, in addition to two novel structures of the FliPQR homologues from *Vibrio mimicus* and *Pseudomonas savastanoi*. The structure reveals a unique topology that presents a loop that wraps around the base of the closed export gate. Mutagenesis studies confirm the crucial role played by the FlhB loop in the export process and suggest potential mechanisms of regulation of opening of the assembly.

## Results

### Conservation of the FliPQR structure

Our previously determined structures of *S.* Typhimurium FliPQR (*7*) and the vT3SS homologue SctRST from *S. flexneri* (*8*) demonstrated that the stoichiometry of the core structure (FliP_5_Q_4_R_1_) is conserved between virulence and flagellar systems. However, classification of the SctRST data revealed variable occupancy of the SctS component (up to a maximum of 4 copies), consistent with our earlier native mass spectrometry measurements (*7*). Furthermore, our native mass spectrometry had also demonstrated that a small proportion of the *P. savastanoi* FliPQR complex contained 5 copies of FliQ, with the predicted 5^th^ FliQ binding site beginning to encroach on the predicted FlhB interaction site. In order to further analyse the structural conservation and stoichiometry of the export apparatus core FliPQR we chose the homologous complexes from the fT3SS of two other bacterial species for structural studies: the *V. mimicus* polar flagellum FliPQR complex, which has a longer FliP sequence including an N-terminal domain conserved in the *Vibrionales* order (Supplementary Fig. 1), and the *P. savastanoi* FliPQR complex, that is a mixture of FliP_5_Q_5_R_1_ and FliP_5_Q_4_R_1_ complexes by native mass spectrometry (*7*). We determined the structures of both complexes using cryo-EM and single particle analysis to 4.1 Å and 3.5 Å respectively (Fig. 1a, Table 1, Supplementary Fig. 2 and Supplementary Fig. 3). Both structures are highly similar to *S.* Typhimurium FliPQR (*7*) (RMSD=1.6 Å over all chains) and *S. flexneri* SctRST (*V. mimicus* FliPQR and SctRST RMSD=1.9 Å and *P. savastanoi* FliPQR and SctRST RMSD=2.3 Å) (*7, 8*).

**Table 1.** Cryo-EM statistics.

|  | EMD-10095, PDB 6S3R | EMD-10096, PDB 6S3S | EMD-10093, PDB 6S3L |
| --- | --- | --- | --- |
| **Data collection and processing** |  |  |  |
| Magnification | 165,000 | 165,000 | 165,000 |
| Voltage (kV) | 300 | 300 | 300 |
| Electron exposure (e<sup>-</sup>/Å<sup>2</sup>) | 48 | 48 | 48 |
| Defocus range (μm) | 0.5–4 | 0.5–4 | 0.5–4 |
| Pixel size (Å) | 0.822 | 0.822 | 0.822 |
| Symmetry imposed | C1 | C1 | C1 |
| Initial particle images (no.) | 503,177 | 1,050,955 | 1,386,230 |
| Final particle images (no.) | 97,987 | 243,489 | 162,408 |
| Map resolution (Å) | 3.5 | 4.1 | 3.2 |
| FSC threshold | 0.143 | 0.143 | 0.143 |
| **Refinement** |  |  |  |
| Initial model used | EMDB-4173 | EMDB-4173 | EMDB-4173 |
| Model resolution (Å) | 3.5 | 4.1 | 3.2 |
| FSC threshold | 0.143 | 0.143 | 0.143 |
| Map sharpening B factor (Å<sup>2</sup>) | -101 | -214 | -97 |
| **Model composition** |  |  |  |
| Non-hydrogen atoms | 12,855 | 12,321 | 13,849 |
| Protein residues | 1,669 | 1,569 | 1,768 |
| Ligands | 0 | 0 | 0 |
| **B factors (Å<sup>2</sup>)** |  |  |  |
| Protein | 36.94 | 101.85 | 43.55 |
| Ligand | N/A | N/A | N/A |
| r.m.s. deviations |  |  |  |
| Bond lengths (Å) | 0.0058 | 0.0066 | 0.01 |
| Bond angles (°) | 0.88 | 0.87 | 0.93 |
| **Validation** |  |  |  |
| MolProbity score | 2.54 | 2.25 | 2.45 |
| Clashscore | 18 | 18.28 | 15.56 |
| Poor rotamers (%) | 2.22 | 0.07 | 2.72 |
| **Ramachandran plot** |  |  |  |
| Favored (%) | 90.82 | 91.86 | 93.64 |
| Allowed (%) | 8.39 | 7.76 | 5.84 |
| Disallowed (%) | 0.79 | 0.39 | 0.52 |

**Fig. 1.**
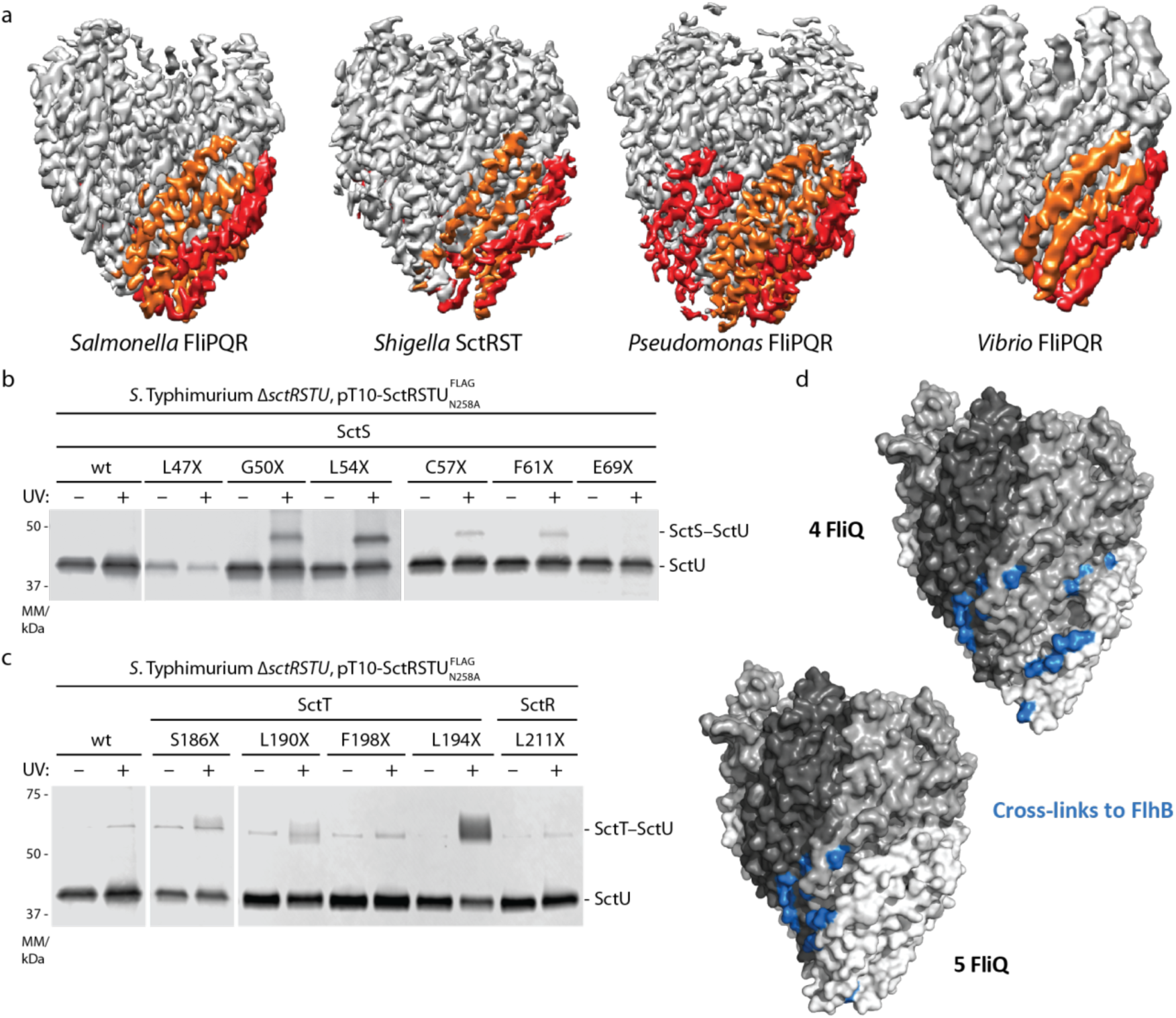
Conservation of the structure of the FliPQR export gate in the closed state. **a**, Cryo-EM volumes calculated in Relion using data from *S.* Typhimurium FliPQR (left, EMD-4173), *S. flexneri* SctRST (centre left, SctR_5_S_4_T_1_ class (*8*) (EMD-4734)), *P. savastanoi* FliPQR (centre right) and *V. mimicus* FliPQR (right). FliQ_2_ and FliQ_4_ are coloured orange and FliQ_1_, FliQ_3_ and FliQ_5_ are coloured red. **b**, Immunodetection of SctUFLAG on Western blots of SDS PAGE-separated crude membrane samples of the indicated *S.* Typhimurium SctS *p*Bpa mutants (denoted with X). Each sample is shown with and without UV-irradiation to induce photocrosslinking of *p*Bpa to neighbouring interaction partners. **c**, As in (b) but testing interactions to SctU with *p*Bpa in SctTand SctR. **d**, Mapping of the confirmed contact points, including those previously identified (*17*) on the structure of FliPQR (*S.* Typhimurium) and a model with a fifth FliQ subunit which is based on the structure of *P. savastanoi* FliPQR.

Consistent with our previous native mass spectrometry data, the structure of *P. savastanoi* revealed an additional FliQ subunit in the complex. In the *S.* Typhimurium and *V. mimicus* FliPQR structures there are four FliP-FliQ units, each the structural equivalent of a FliR subunit (*7*), but the fifth FliP is missing a FliQ. In the *P. savastanoi* structure, FliQ_5_ binds the remaining FliP subunit in the same way as in the other FliP-FliQ units. This FliQ_5_ subunit is located close to the site on FliPQR we previously identified as important for interaction with FlhB/SctU (*7, 17*). Mapping of more a extensive *in vivo* photocrosslinking analysis based on covariance (*16*) between SctU and SctR, SctS, and SctT supports a binding site for SctU that involves large parts of helix 2 of SctS and helix 4 of SctT (Fig. 1b-c and Supplementary Fig. 4). Mapping of the residues on the structure of FliPQR and a model of FliPQR containing a fifth FliQ subunit reveal a large binding site in the complex containing four FliQ subunits and a more compact binding site when a fifth FliQ subunit is modelled (Fig. 1d). In this way FliQ_5_/SctS_5_ might be compatible with FlhB/SctU binding, depending on the unknown structure of the FlhB/SctU transmembrane domain (FlhB/SctU_TM_), but addition of a sixth Fli/SctQ using the same helical parameters, would block this site.

### Architecture of the FliPQR-FlhB export gate complex

We have observed four FliQ subunits in the *S.* Typhimurium and *V. mimicus* FliPQR and the *S. flexneri* SctRST structures but as we have previously observed FliQ to be sensitive to dissociation by detergent in the purification process (*7, 8*), it was possible that the 5^th^ FliQ is a genuine component of the complex but was lost in the purification of less stable homologues. As the stoichiometry of FliQ has potentially large implications for the placement of FlhB in the system (Fig. 1d), we endeavoured to produce the more physiologically relevant FliPQR-FlhB complex.

After extensive screening of detergents, constructs with different placement of affinity tags and sequences from a variety of species for co-expression and co-purification of FlhB with FliPQR, we were able to prepare a monodisperse sample of the complex from *V. mimicus* (Fig. 2a). We analysed this sample by cryo-EM and determined the 3.2 Å structure of the complex (Fig. 2b, Table 1 and Supplementary Fig. 5), revealing a single copy of FlhB added to the previously observed FliP_5_Q_4_R_1_ complex. The FliPQR subcomplex in this structure is very similar to the structure of the FliPQR complex in the absence of FlhB (RMSD=0.6 Å), while FlhB is observed to contain four long helices in the putative TM domain (FlhB_TM_), forming two distinct hairpins that are wrapped around the outside of the FliPQR complex. This extensive interaction surface between FlhB and FliPQR reveals FlhB to be an integral part of the core of the export apparatus instead of an accessory factor. The opened out structure of the 4 predicted TM helices of the FlhB_TM_ domain once again highlights the potential dangers in predicting complex structures in the absence of some of the subunits. The soluble, globular, cytoplasmic domain (FlhB_C_) is not visible, likely due to flexibility in the linker between the two domains. We tested whether this disorder of FlhB_C_ relative to FliPQR-FlhB_TM_ is due to the presence of the detergent micelle in our sample by imaging the sample in the amphipol A8-35, perhaps a better mimic of the proteinaceous environment relevant to the assembled T3SS (*7*), but we did not observe any additional density resulting from ordering of FlhB_C_ (data not shown).

**Fig. 2.**
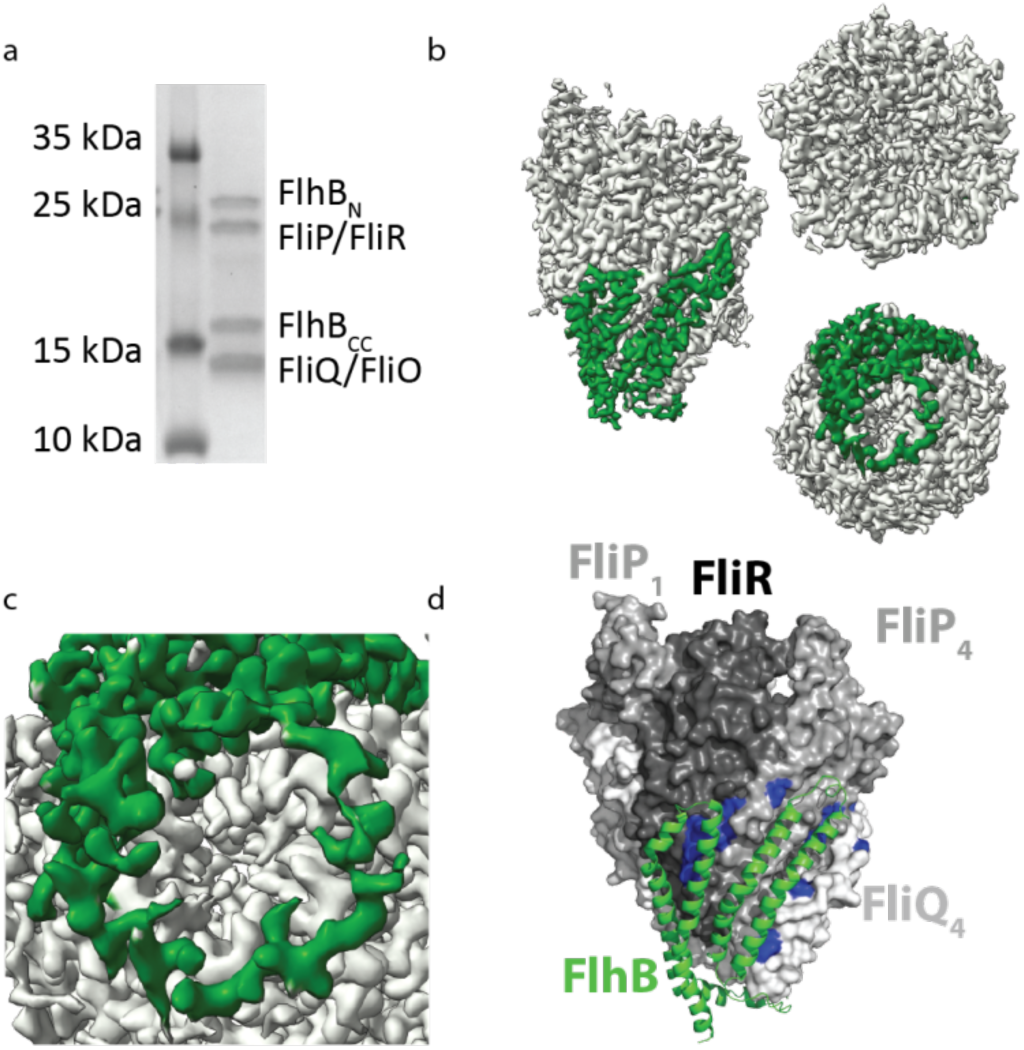
Architecture of the FliPQR-FlhB export gate complex. **a**, SDS-PAGE gel of the *V. mimicus* FliPQR-FlhB complex. **b**, Cryo-EM density map of the FliPQR-FlhB complex with the density corresponding to FlhB coloured in green. **c**, Zoom of the FlhB loop at the bottom of the complex. **d**, Model of FliPQR (surface) and FlhB (cartoon, green) with residues in FliP/SctR and FliR/SctT cross-linking to FlhB/SctU highlighted in blue as in Fig. 1d.

Intriguingly, the two helical hairpins of FlhB are joined by a loop (FlhB_L_) that literally loops around the (closed) entrance of the FliPQR gate (Fig. 2c). Consistent with our previous prediction (*7*) and crosslinking analysis (Fig. 1), FlhB contacts the site across FliP_5_ and FliR, but it additionally contacts cross-linkable residues in the FliP_4_ subunit (Fig. 2d). As previously predicted (*7*), hydrophobic cavities between FliP and FliQ, in addition to lateral cavities between the FlhB hairpins and the FlhB/FliQ interface, are observed to contain densities consistent with lipid or detergent molecules although these could not be modelled unambiguously in the current volume (Supplementary Fig. 6).

### Structure of the hydrophobic domain of FlhB

The density corresponding to FlhB was of sufficient quality to build an atomic model of the structure using only sequence information (Fig. 3a and Supplementary Fig. 5). The topology of FlhB_TM_ is unusual; the helices 1 and 4 neighbour each other in the middle of the structure, while helix 2 and 3 flank the central pair on either side (Fig. 3b,c). In order to further validate the topology of FlhB we compared our model to contacts derived from evolutionary co-variation (Fig. 3d and Supplementary Fig. 7). This shows strong contacts between helices 1 and 2, 3 and 4 and 1 and 4 but an absence of contacts between helices 2 and 3, which is inconsistent with a helical bundle, but consistent with our more extended and topologically unusual structure.

**Fig. 3.**
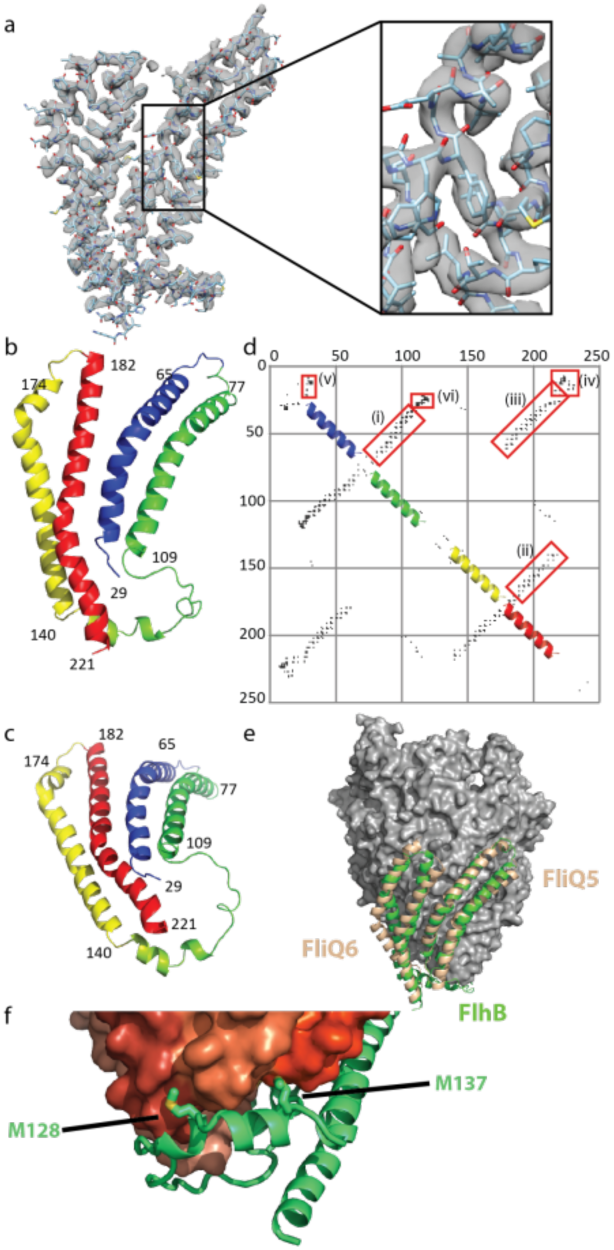
Structural analysis of the FlhB hydrophobic domain. **a**, Quality of the cryo-EM volume corresponding to FlhB. Zoom box shows the fit to density of the model. **b**,**c**, Rainbow colouring of the FlhB model with numbers indicating the N and C-termini of the four helices (*V. mimicus* numbering). **d**, Evolutionary co-variation within FlhB calculated using RaptorX (*18*). Only contacts with a probability greater than 0.5 are plotted. Red boxes highlight the interaction between helix 1 and 2 (i), helix 1 and 4 (ii), helix 3 and 4 (iii), the N-terminus and FlhB_CN_ (iv), within the N-terminus (v) and between FlhB_L_ and the N-terminus (vi). **e**, Overlay of FlhB (green) and a modelled FliQ_5_ and FliQ_6_ following the same helical parameters as FliQ_1_ to FliQ_4_ in *V. mimicus*. **f**, Zoomed view of the interaction between the FlhB loop and FliQ, highlighting the intercalation of conserved hydrophobic residues in FlhB between the FliQ subunits.

Despite observing up to five FliQ subunits in FliPQR structures, there are only four FliQ molecules in this structure. In fact, the hairpin composed of FlhB helices 1 and 2 is bound to the site occupied by FliQ_5_ in our *P. savastanoi* FliPQR structure, packing on FliP_5_, whilst helices 3 and 4 pack on FliR. Thus the presence of FliQ_5_ would block binding of FlhB (Fig. 3e), suggesting that the fifth FliQ binds to the complex in a non-native fashion due to the absence of FlhB in the overexpression system. This superposition of FlhB and FliQ_5_ also reveals that the hairpin of helices 1 and 2 of FlhB adopts a very similar structure to FliQ despite the fact they are topologically distinct, with helix 2 of FlhB being structurally equivalent to helix 1 of FliQ and vice versa, i.e. the directionality of the hairpin is reversed along the long axis. Modelling FliQ_5_ and FliQ_6_ using the same helical parameters by which FliQ_1_ to FliQ_4_ are related reveals that FlhB continues the spiral of FliQ subunits and even helices 3 and 4 follow the same parameters (Fig. 3e) despite not interacting with a FliP subunit. Given the very different topologies of the two proteins, the level to which FlhB helices 1 and 2 and FliQ superpose is surprising. An evolutionary relationship between FlhB and FliQ is unlikely due to the topology differences, suggesting that the similarity of the structures is a result of convergent evolution and the need to form this helical assembly.

### An extended loop between helices is essential for secretion

The unusual topology of FlhB_TM_ means that a long loop, FlhB_L_, between helix 2 and 3 (residues 110-139) connects the two hairpins of the structure. Most unexpectedly, this loop, c which contains the most highly conserved residues within FlhB (Supplementary Fig. 8), is seen to wrap around the base of the PQR complex, contacting each of the FliQ subunits in turn and inserting conserved hydrophobic residues into the cavities between the FliQs (Fig. 3f). The loop structure also reveals how a single FliQ residue can co-evolve with multiple FlhB_L_ residues. Although FlhB_L_ in isolation doesn’t further constrict the base of the already closed PQR complex, the aperture does become significantly smaller when taking into account the poorly resolved termini of FlhB_TM_ (Supplementary Fig. 9). Therefore FlhB_L_ could contribute to export gate closure via trapping of the FlhB N-terminus and the linker connecting FlhB_TM_ to FlhB_C_, in the direct line of the export pathway or by pinning the FliQ subunits closed. A mutation in the FlhB N-terminus had been reported (*19*) to act as a Δ*fliHI* bypass mutant (the ATPase and its regulator), presumed to be involved in controlling the opening of the export channel. In the FliPQR-FlhB complex the equivalent residue (FlhB_P28_ in *S.* Typhimurium, FlhB_A28_ in *V. mimicus*) locates very close to the pore entrance (Supplementary Fig. 9). In the *S.* Typhimurium SctRSTU complex the corresponding residue strongly photocrosslinks to SctS (Supplementary Fig. 4), supporting the notion that SctU mediates gating of the export apparatus core complex.

We decided to further probe the function of FlhB_L_ using mutagenesis in the motile *E. coli* strain W (Fig. 4a and Supplementary Fig. 10). Given that opening of the FliPQR-FlhB aperture would require a conformational change in FlhB_L_, we hypothesised two mechanisms for FlhB_L_ such conformational changes. FlhB_L_ could either move away from the entrance to the gate through a hinging motion like a lid, or it could extend into a structure with less secondary structure in order to stay in contact with the binding sites on the opening FliQ subunits, reminiscent of a sphincter.

**Fig. 4.**
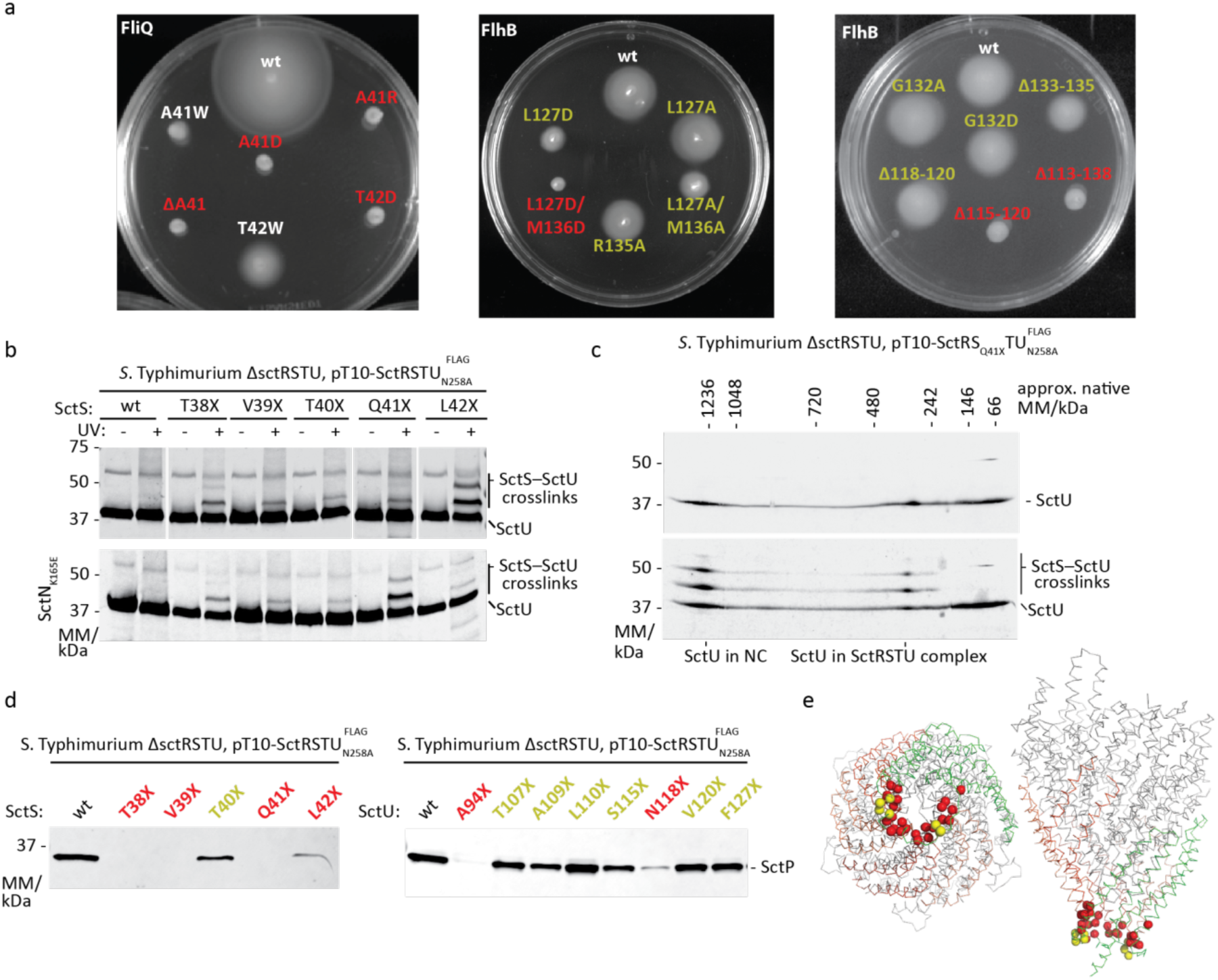
Functional analysis of FlhB_L_. **a**, Motility in soft agar of *E. coli* W ΔFliOPQR complemented with plasmids expressing FliOPQR with the indicated mutations in FliQ (left) and *E. coli* W ΔFlhB complemented with plasmids expressing FlhB with the indicated mutations (*E. coli* numbering) (right). **b**, Immunodetection of SctU^FLAG^ on Western blots of SDS-Page separated crude membrane samples of the indicated *S.* Typhimurium SctS *p*Bpa mutants (denoted with X). Each sample is shown with and without UV-irradiation to induce photocrosslinking of *p*Bpa to neighbouring interaction partners. Crosslinks between SctS_*p*Bpa_ and SctU^FLAG^ are indicated. Crosslinking analysis was performed in the wild type and in an ATP hydrolysis-deficient SctN_K165E_ mutant that is incapable of type III secretion. **c**, Immunodetection of SctU^FLAG^ on Western blots of 2D blue native/SDS-PAGE separated crude membrane samples of the indicated *S.* Typhimurium SctS *p*Bpa mutant. The sample is shown with and without UV-irradiation to induce photocrosslinking of *p*Bpa to neighbouring interaction partners. Crosslinks between SctS_*p*Bpa_ and SctU^FLAG^ in the SctRSTU assembly intermediate and in the assembled needle complex (NC) are indicated. **d**, Immunodetection of the early T3SS substrate SctP on Western blots of SDS-PAGE separated culture supernatants of the indicated *S.* Typhimurium SctS *p*Bpa mutants. **e**, Structure of FliPQR-FlhB highlighting mutation sites that impaired motility or secretion in red and mutation sites that had no or only a small effect in yellow.

Mutations in either the conserved hydrophobic residues of FlhB_L_ that insert between the FliQ subunits (Fig. 3f) or the highly conserved loop of FliQ severely reduced motility (Fig. 4a) without affecting binding of FlhB to FliPQR (Supplementary Fig. 11). Although substitution with the bulky, hydrophobic amino acid tryptophan and removal of bulky sidechains only reduced motility, introduction of charged residues completely abolished motility, suggesting that secretion can proceed at lower efficiency when the FliQ-FlhB_L_ interaction is only reduced rather than completely disrupted as in the aspartate mutations.

We performed an extensive *in vivo* photocrosslinking analysis to validate the interactions and functional relevance of the corresponding SctU_L_ in the vT3SS-1 of *S.* Typhimurium. While no crosslinks to SctS could be identified with the artificial crosslinking amino acid *p*-benzoyl-phenylalanine (*p*Bpa) introduced into SctU_L_ itself (Supplementary Fig. 12), numerous crosslinks were identified with *p*Bpa in the lower part of SctS that faces SctU_L_ (Fig. 4b). Using 2-dimensional blue native/SDS PAGE, we could show that the crosslink observed with SctS_Q41X_ occurred not only in the SctRSTU assembly intermediate but also in the assembled needle complex (Fig. 4c), adding further support to the idea that the structure of the isolated complex represents the structure of the complex in the full assembly. The observed crosslinks were independent of functional secretion of the vT3SS, indicating that assembly of needle adapter, the inner rod, and needle filament does not lead to a conformational change of this part of the SctRSTU complex (Fig. 4b). Strikingly, introduction of *p*Bpa at several positions of SctS led to a strong defect in secretion but not SctS-SctU_L_ interaction, highlighting the relevance of this site for secretion function of T3SS (Fig. 4d), while *p*Bpa substitutions within SctU_L_ were more functionally neutral. In total, we found a large number of residues at the FliQ/SctS-FlhB_L_/SctU_L_ interface that are required for type three secretion (Fig. 4e).

Strong cross-linking between SctS and SctU even in the assembled needle complex and loss of function in more disruptive mutations in which the interaction between FliQ and FlhB_L_ is altered through the introduction of charged residues suggest that this interaction is important for activity and FlhB_L_ is not simply one of the closure points of the complex in assembly intermediates. If this interaction is maintained in the open state of the export gate, a more extended conformation of FlhB_L_ would be required. Consistent with this idea, deletions of six or more residues in FlhB_L_ led to loss of motility (Fig. 4a).

We further investigated the function of FlhB_L_ through more targeted mutations. FlhB_L_ is a largely extended polypeptide with little secondary structure, but a short stretch at its C-terminus is helical. Interestingly, a mutation of a glycine in this α-helix in the FlhB homologue YscU disrupted secretion in the *Yersinia* vT3SS (*20*). The equivalent mutation in *E. coli* FlhB and mutation of the conserved positively charged residue R135 (*E. coli* W numbering) had little effect (Fig. 4a) assessed at the level of motility although we cannot rule out more subtle effects on the efficiency with which secretion occurs.

## Discussion

In this study we show that FlhB_TM_ is part of the export gate complex together with FliPQR. Two pairs of helices of FlhB bind to FliPR through a structure mimicking the shape of FliQ, despite topological reversal, an example of molecular convergent evolution. The unusual topology of FlhB places helices 2 and 3 apart from each other allowing them to mount a loop, FlhB_L_, onto the cytoplasm-facing surface of the export gate. Although the way in which FlhB_L_ wraps around the closed pore suggests a role in maintaining the closed state, our structures of FliPQR/SctRST in the absence of FlhB/SctU are also closed (*7, 8*), as is the complex in the context of the assembled T3SS (*21*), suggesting instead that FlhB may be involved in opening of the gate rather than locking it closed. We further propose that in the process of opening FlhB_L_ forms a more extended structure, implying that it undergoes cycles of extension and contraction as the export gate opens and closes.

The location and the topology of FlhB_TM_ place the N-terminus, FlhB_CN_ and FlhB_L_ in close proximity just underneath the aperture of the gate. Although the resolution of the map is poor in the region of the cytoplasmic face of the complex, it is possible to trace the approximate position of the FlhB N-terminus and FlhBCN (Supplementary Fig. 9). The close association of the N-terminus and FlhB_CN_ is consistent with the strong contacts derived from evolutionary co-variation between the N-terminus and the N-terminal part of the cytoplasmic domain (FlhB_CN_) (Fig. 3c) and a genetic interaction in *S.* Typhimurium between E11 and E230, in the N-terminus of FlhB and FlhB_CN_ respectively (*22*). Furthermore, the direction in which FlhB_CN_ leaves the export gate implies that, in the context of the assembled T3SS nanomachine, FlhB_C_ could be located anywhere between the FliPQR-FlhB gate and the nonameric ring of the FlhA cytoplasmic domain below (Fig. 5a). It is conceivable that the conformational changes required for opening of the export gate are propagated via pulling forces imparted on helix 4 of FlhB_TM_, which is linked to the other helices of FlhB_TM_ via a conserved network of buried charged residues (Fig. 5b) including D208 that has been demonstrated to play a role in motility (*23*). Such forces could initiate in the cytoplasmic domain of FlhA, which binds to substrate-chaperone complexes and has been demonstrated to exist in multiple conformations (*24, 25*) marking it as the only component of the export apparatus observed in multiple conformational states to date. As FlhB_C_ is thought to interact with substrates just before they pass through the export gate (*26*), it is possible that transition of FlhA_C_ between the different conformations pulls on FlhB_C_, either directly or via substrate, thereby pulling on the FlhB_TM_ network. Alternatively, changes in the FlhA_TM_ domain that are triggered via FlhA_C_ or by the proton-motive force (*27*) could directly influence the conformation of the FlhB_TM_ domain.

**Fig. 5.**
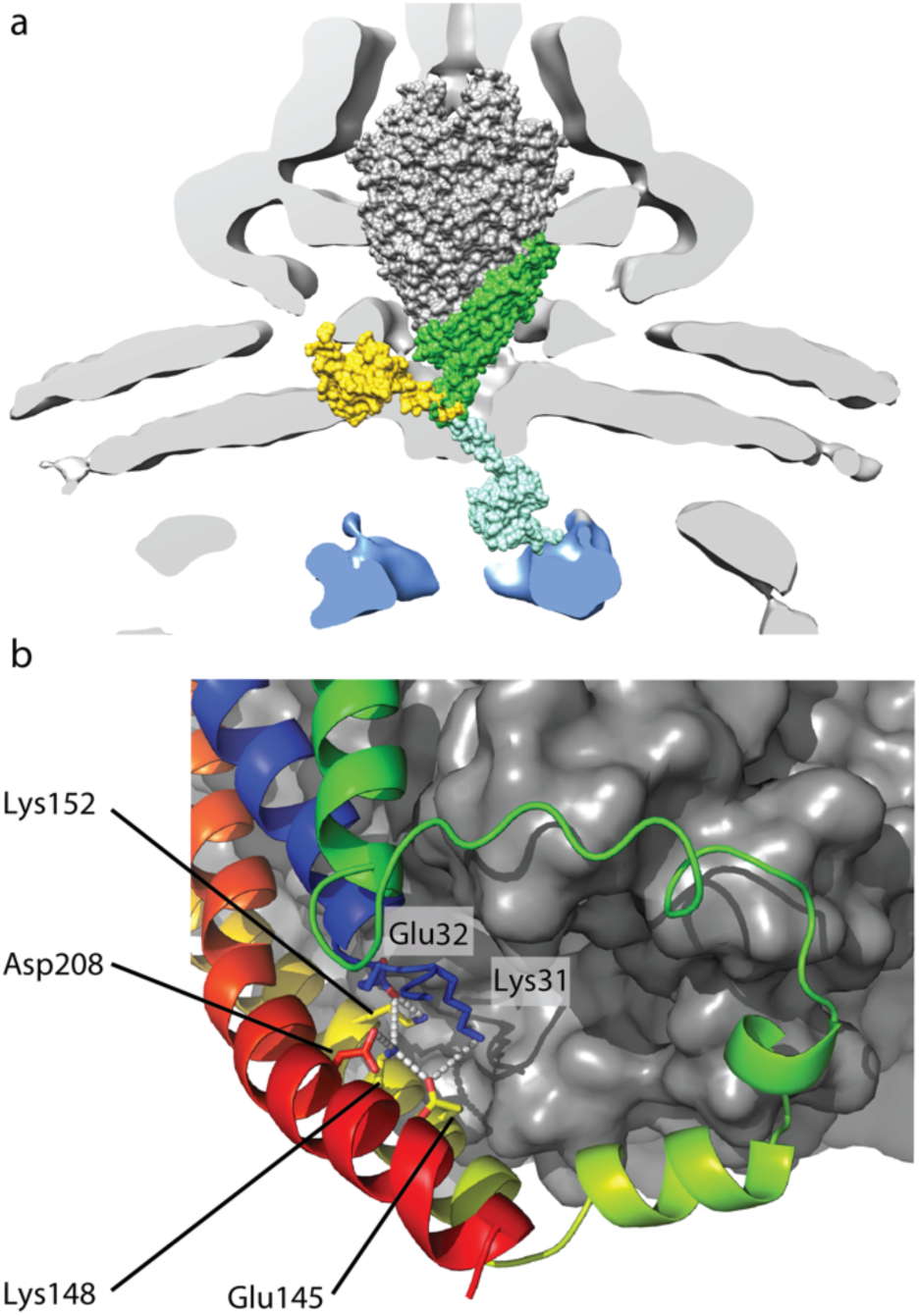
Position of FlhB_C_ in the complete T3SS. **a**, Placement of FliPQR-FlhB in a tomographic reconstruction of the *Salmonella* SPI-1 vT3SS (EMD-8544) (*30*). FliPQR is shown in grey, FlhB_TM_ in green, two possible positions of FlhB_C_ (PDB: 3b0z) (*13*) in yellow and light blue and the density corresponding to the FlhA homologue is highlighted in blue. **b**, Network of salt bridges formed by conserved charged residues in FlhB (*V. mimicus* numbering).

The mechanism of suppressor mutations in the N-terminal residues of FlhB, such as the P28T mutation that rescues motility of a strain deleted for the ATPase complex (Δ*fliHI)* (*19, 28*), has long been mysterious. Our structure, demonstrating the clustering of the N-terminus, FlhB_CN_ and FlhB_L_ at the cytoplasmic entrance of the gate, suggests that they may rescue function by altering the dynamics of the closure point in order to facilitate opening of the export gate. This notion is further supported by the interaction we observe of SctU_F24*p*Bpa_ (equivalent to P28T) with SctS. This direct functional link between the ATPase and the gating mechanism, in conjunction with a host of other mutational data in FlhA and FlhB, suggests that cycles of ATP hydrolysis may induce conformational changes in the export gate, presumably via the FliJ stalk of the ATPase complex interacting with the FlhA ring that is positioned between the ATPase and the export gate.

Finally, FlhB/SctU is known to play a key role in substrate switching, an event which requires autocatalytic cleavage of the NPTH sequence in FlhB_C_/SctU_C_ (*14, 15, 26, 29*). Although we do not observe the residues involved in switching in our export gate structure, the fact that we are able to cross-link SctU in fully assembled basal bodies, using the same residues as in the purified complex, suggests that the gating mechanism discussed here is likely applicable regardless of substrate. Clearly future studies will need to focus on observing gating and switching events.

In summary, our structure of FlhB as part of the export gate complex extends our understanding of the regulation of the T3SS export apparatus and suggests possible mechanisms of export gate opening.

## Methods

### Materials

Chemicals were from Sigma-Aldrich unless otherwise specified. The detergents n-dodecylmaltoside (DDM) and lauryl maltose neopentyl glycol (LMNG) and the amphipol A8-35 were from Anatrace. *p*-benzoyl-phenylalanine was from Bachem. Primers are listed in Supplementary Table 2 and were synthetized by Invitrogen or Eurofins.

### Bacterial strains and plasmids

Bacterial strains and plasmids used in this study are listed in Supplementary Table 3. Plasmids were generated by Gibson assembly of PCR fragments using the NEBuilder HiFi Master Mix (NEB) or in vivo assembly (*31*). Fragments were created by PCR with the relevant primers using Q5 polymerase (NEB) and genomic DNA templates obtained from DSMZ (*Vibrio mimicus* strain DSM 19130 and *Pseudomonas savastanoi*, pv. phaseolicola 1448A strain DSM 21482). Gibson assembly and PCR were carried out following the manufacturer’s recommendations. *Escherichia coli* W for motility assays was obtained from DSMZ (DSM 1116). Bacterial cultures were supplemented as required with ampicillin (100 μg/mL) or kanamycin (30 μg/mL or 60 μg/mL for large scale expression in TB medium). *S*. Typhimurium strains were cultured with low aeration at 37°C in Luria Bertani (LB) broth supplemented with 0.3 M NaCl to induce expression of genes of SPI-1. As required, bacterial cultures were supplemented with tetracycline (12.5 µg/ml), streptomycin (50 µg/ml), chloramphenicol (10 µg/ml), ampicillin (100 µg/ml) or kanamycin (25 µg/ml). Low-copy plasmid-based expression of SctRSTU^FLAG^ was induced by the addition of 500 µM rhamnose to the culture medium.

### Generation of chromosomal deletion mutants

Electrocompetent *E. coli* W expressing λ Red recombinase from plasmid pKD46 were transformed with DNA fragments containing a chloramphenicol resistance cassette surrounded by sequences homologous to the gene of interest as described in Supplementary Table 3. Colonies were selected on LB agar containing chloramphenicol (20 μg/mL) and transformed again with pCP20 and grown on LB agar containing ampicillin (100 μg/mL) at 30 °C. Finally, clones were grown in LB media at 37 °C. Deletion mutations were confirmed by PCR. All *Salmonella* strains were derived from *Salmonella enterica* serovar Typhimurium strain SL1344 (Hoiseth and Stocker, 1981) and created by allelic exchange as previously described (Kaniga *et al.*, 1994).

### Purification of export gate complexes

FliOPQR or FliOPQR-FlhB were expressed in *Escherichia coli* BL21 (DE3) as a single operon from a pT12 vector (Supplementary Table 3) as described previously (*7*). Briefly, cells were grown at 37 °C in TB media containing rhamnose monohydrate (0.1%), harvested by spinning at 4,000 g, resuspended in TBS (100 mM Tris, 150 mM NaCl, 1 mM EDTA, pH 8) and lysed in an EmulsiFlex C5 homogenizer (Avestin). Membranes were prepared from the cleared lysate by ultracentrifugation at 40,000 rpm in a 45 Ti rotor (Beckmann) for 3 hours. Membranes were solubilized in 1% (w/v) LMNG and proteins were purified using a StrepTrap column (GE Healthcare). The resin was washed in TBS containing 0.01% (w/v) LMNG and proteins were eluted in the wash buffer supplemented with 10 mM desthiobiotin. Intact complexes were separated from aggregate by size-exclusion chromatography in TBS containing 0.01% (w/v) LMNG (S200 10/300 increase or Superose 6 increase, GE Healthcare).

For preparation of FliPQR-FlhB solubilised by the amphipol A8-35, the protein was purified in DDM using 1% (w/v) for extraction from the membrane and 0.02% (w/v) subsequently. Eluate from the StrepTrap column was mixed with amphipol at a ratio of amphipol to protein of 10:1. After incubating for one hour, the sample was dialysed into TBS using a 10,000 MWCO Slide-A-Lyzer device (ThermoFisher Scientific) overnight followed by size-exclusion chromatography on a Superose 6 increase column using TBS as the running buffer.

### Sample preparation for cryo-EM

3 μl of purified complex at 1 to 3.6 mg/ml were applied to glow-discharged holey carbon-coated grids (Quantifoil 300 mesh, Au R1.2/1.3). Grids were blotted for 3 s at 100% humidity at 22 °C and frozen in liquid ethane using a Vitrobot Mark IV (FEI). For samples solubilised in detergent, blotting was preceded by a wait time of 5 to 10 seconds. *V. mimicus* FliPQR was supplemented with 0 mM, 0.05 mM, 0.5 mM or 3 mM fluorinated Fos-Choline prior to grid preparation.

### EM data acquisition and model building

All data contributing to the final models were collected on a Titan Krios (FEI) operating at 300 kV. All movies were recorded on a K2 Summit detector (Gatan) in counting mode at a sampling of 0.822Å/pixel, 2.4 e– Å^−2^/frame, 8 s exposure, total dose 48 e–/ Å^−2^,20 fractions written. Motion correction and dose weighting were performed using MotionCor implemented in Relion 3.0 (*32*) (*V. mimicus* FliPQR-FlhB and *P. savastanoi* FliPQR) or using Simple-unblur (*33*) (*V. mimicus* FliPQR). CTFs were calculated using CTFFIND4 (*34*). Particles were picked in Simple and processed in Relion 2.0 (*35*) and 3.0 (*32*) as described in Supplementary Fig. 2, Supplementary Fig. 3 and Supplementary Fig. 5. Atomic models of FliPQR and FlhB were built using Coot (*36*) and refined in Phenix (*37*).

### Motility assays

*E. coli* W strains WL1 or WL2 (Supplementary Table 3) were transformed with plasmids encoding FliOPQR or FlhB containing the mutations to be tested. 3 μl of a saturated overnight culture were injected into soft agar plates (0.28% agar, 2YT media, containing ampicillin (100 μg/mL) or kanamycin (30 μg/mL) and 0.1% arabinose or 0.5% rhamnose monohydrate as appropriate) and incubated at room temperature.

### *S*. Typhimurium *vT3SS secretion assay*

Proteins secreted into the culture medium via the vT3SS-1 were analyzed as described previously (Monjarás Feria *et al.*, 2015). *S*. Typhimurium strains were cultured with low aeration in LB broth supplemented with 0.3 M NaCl at 37°C for 5 h. Bacterial suspensions were centrifuged at 10,000 x *g* for 2 min and 4°C to separate whole cells and supernatants. Whole cells were resuspended in SDS PAGE loading buffer. Supernatants were passed through 0.2 µm pore size filters and supplemented with 0.1 % Na-desoxycholic acid. Proteins in the supernatant were precipitated with 10 % tricholoacetic acid (TCA) for 30 min at 4°C and pelleted via centrifugation at 20,000 x *g* for 20 min and 4°C. Pellets containing precipitated proteins were washed with acetone and resuspended in SDS PAGE loading buffer. Whole cell samples and secreted proteins were analyzed by SDS PAGE, Western blotting, and immunodetection.

### In vivo photocrosslinking

*In vivo* photocrosslinking was carried out as described previously (Farrell *et al.*, 2005; Dietsche *et al.*, 2016) with minor modifications. In order to enhance expression of vT3SS-1, *S*. Typhimurium strains expressed HilA, the master transcriptional regulator of SPI-1 T3SS, from a high copy plasmid under the control of an arabinose-inducible *P*_araBAD_ promoter (Guzman *et al.*, 1995). Bacterial cultures were grown in LB broth supplemented with, 0.3 M NaCl, 1 mM *p*Bpa and 0.05 % arabinose at 37°C for 5 h. 5 ODU of bacterial cells were harvested and washed once with 5 ml chilled PBS to remove residual media. Bacteria were pelleted by centrifugation at 4,000 x *g* for 3 min and 4°C and afterwards resuspended in 1 ml of chilled PBS. Bacterial suspensions were transferred into 6-well cell culture dishes and irradiated for 30 min with UV light (λ=365 nm) on a UV transilluminator table (UVP). Subsequently, bacterial cells were pelleted by centrifugation at 10,000 x *g* for 2 min and 4°C. Samples were stored at −20°C until use.

### Crude membrane preparation

Crude membranes were purified following the published protocol (Dietsche *et al.*, 2016). 5 ODU of bacterial cells were resuspended in 750 µl lysis buffer (50 mM triethanolamine pH 7.5, 250 mM sucrose, 1 mM EDTA, 1 mM MgCl2, 10 µg/ml DNAse, 2 µg/ml lysozyme, 1:100 protease inhibitor cocktail), and incubated on ice for 30 minutes. Cell slurries were lysed via continuous bead milling. Intact cells, beads and debris were removed by centrifugation for 10 min at 10,000 x *g* and 4°C. Supernatants were centrifuged for 50 min at 52,000 rpm and 4°C in a Beckman TLA 55 rotor to pellet bacterial membranes. Pellets containing crude membranes were store at −20°C until use. Samples were analyzed by SDS PAGE, Western blotting, and immunodetection.

### Western blotting and immunodetection

Samples were loaded onto SERVAGel^™^ TG PRiME 8-16% precast gels and transferred on PVDF membranes (Bio-RAD). Proteins were detected with primary antibodies anti-*St*_1_SctP (InvJ) (Monjarás Feria *et al.*, 2015) (1:2000) or M2 anti-FLAG (1:10,000) (Sigma-Aldrich). Secondary antibodies (Thermo-Fisher) were goat anti-mouse IgG Dylight 800 conjugate (1:5000). Scanning of the PVDF membranes and image analysis was performed with a Li-Cor Odyssey system and Image Studio 3.1 (Li-Cor).

## Acknowledgements

We thank E. Johnson and A. Costin of the Central Oxford Structural Microscopy and Imaging Centre for assistance with data collection. H. Elmlund (Monash) is thanked for assistance with access to SIMPLE code ahead of release. The Central Oxford Structural Microscopy and Imaging Centre is supported by the Wellcome Trust (201536), the EPA Cephalosporin Trust, the Wolfson Foundation, and a Royal Society/Wolfson Foundation Laboratory Refurbishment Grant (WL160052). Work performed in the lab of S. M. Lea was supported by a Wellcome Trust Investigator Award (100298) and an MRC Program Grant (M011984). L.K. is a Wellcome Trust PhD student (109136).

## Author Contributions

L.K. performed experiments, did strain and plasmid construction, complex purification, native mass spectrometry, cryo-EM grid optimization, cryo-EM data analysis, and model building and analysis. J.D. performed experiments, cryo-EM grid optimization, and data collection. J.F. performed experiments, complex purification. Work in the laboratory of SW related to this project was funded by the German Center for Infection Research (DZIF), project TTU06.801/808. A.Z., S.B., and R.D. generated pBpa mutants, performed crosslinking experiments and secretion assays, and analyzed data. S.W. designed injectisome functional experiments, analyzed data S.J. and S.M.L. supervised experimental work and wrote the first draft of the paper with L.K. All authors contributed to and commented on the final manuscript.

