## Supplementary Figures and Tables for "The flagellar substrate specificity switch protein FlhB assembles onto the extra-membrane export gate to regulate type three secretion"

**Supplementary Fig. 1 Alignment of FliP sequences.**

```
Salmonella_enterica          --------------------MRRLLFLSLAGLWLFSPAAA--------------------      0
Photobacterium_angustum      -----------MLKNKFTLITWMAFLAISVLLSSINIAHA---------VEADVVNSTTA      41
Aliivibrio_fischeri          ---------------MLKTNFIAAVLAAFSLFFMASPAWAVDDLASSIPQTAASAEGMTV      46
Vibrio_mimicus               ------MKRTQRLNLTSWLTTGLLTWLLSGMLGFASLAFAEEPLNTGIPSTAAGASSVTV      55
Vibrio_alginolyticus         ----------MTKGNPIRLLLSTLLLFVGVLFSSFTFAEA---EESALPDNLADGSSVTV      48

Salmonella_enterica          ---------------------AQLPGLISQPLAGGGQSWSLSVQTLVFITSLTFLPAILL      59
Photobacterium_angustum      TAMSAETT---SGHLQAARNGGGIPALSVTVNPDGSEDYSVTLQILALMTALGFLPAMVI      97
Aliivibrio_fischeri          TAMEKEAGAAKTIAKGS-SSGSGIPAFTMTTNANGGEDYSVNLQILGLMTMLGFLPAMVI      104
Vibrio_mimicus               TALEKEQGNAKTIALGSSSGGSGIPAFTMTTNPDGSEDYSINLQILALMTMLGFLPAMVI      114
Vibrio_alginolyticus         QAMQSDTGASRAM---SVSSGGGIPAFTMTTNPDGSEDYSVTLQILALMTMLGFLPAMVI      104
                              :              :   .. :*.:      .*.:.:*:.:* * ::* * ****:::

Salmonella_enterica          MMTSFTRIIIVFGLLRNALGTPSAPPNQVLLGLALFLTFFIMSPVIDKIYVDAYQPFSEQ      119
Photobacterium_angustum      LMTSFTRIVVVMSILRQAMGLQQTPSNQVIIGIALFLTFFVMSPVIDKINANAVQPYINE      157
Aliivibrio_fischeri          LMTSFTRIVVVMSILRQAMGLQQTPSNQVIIGIAMFLTFFIMAPVFDKVNNTAIQPYINE      164
Vibrio_mimicus               LMTSFTRIVVVMSILRQAMGLQQTPSNQVIIGIALFLTFFIMAPVFNQINEQAVQPYLNE      174
Vibrio_alginolyticus         LMTSFTRIVVVMSILRQAMGLQQTPSNQVIIGIALFLTFFVMSPVLNEINDKAVQPYLNE      164
                             :*******::*:.:**:*:*  .:* ***::*:*:*****:*:**::::   * **: ::

Salmonella_enterica          KISMQEALDKGAQPLRAFMLRQTREADLALFARLANSGPLQGPEAVPMRILLPAYVTSEL      179
Photobacterium_angustum      QITAKEALAKAEDPMRQFMLKQTRVKDLETFVNMSGS-TADDPQTVPLTVLIPAFITSEL      216
Aliivibrio_fischeri          EITAREAFDVAQVPMKEFMLAQTRVKDLETFVNMSGV-EAENPEDVPITVVIPAFITSEL      223
Vibrio_mimicus               QISARQAFDLAQEPMKAFMLKQTRIKDLETFVEMSGS-QVTAPEQVSMAVLIPAFITSEL      233
Vibrio_alginolyticus         QVTAREAFDAAQAPMKAFMLKQTRIKDLETFVTMSGE-QVENPEDVSMAVLIPAFITSEL      223
                             ::: ::*:  .  *:: *** ***  **  *. ::.      *: * : :::**::****

Salmonella_enterica          KTAFQIGFTIFIPFLIIDLVIASVLMALGMMMVPPATIALPFKLMLFVLVDGWQLLMGSL      239
Photobacterium_angustum      KTAFQIGFMLFLPFLIIDLVVASILMAMGMMMLSPMIVSLPFKLMLFVLVDGWNLILSTL      276
Aliivibrio_fischeri          KTAFQIGFMLFLPFLIIDLVVASVLMAMGMMMLSPMIVSLPFKLMLFVLVDGWNLILSTL      283
Vibrio_mimicus               KTAFQIGFMLFLPFLIIDLVVASVLMAMGMMMLSPMIVSLPFKLMLFVLVDGWNLILSTL      293
Vibrio_alginolyticus         KTAFQIGFMLFLPFLIIDLVVASVLMAMGMMMLSPMIVSLPFKLMLFVLVDGWNLILSTL      283
                             ******** :*:********:**:***:****: *  ::**************:*::.:*

Salmonella_enterica          AQSFYS      245
Photobacterium_angustum      AGSFGT      282
Aliivibrio_fischeri          AGSFAT      289
Vibrio_mimicus               AGSFAL      299
Vibrio_alginolyticus         AGSFAL      289
                             * **
```

Sequence alignment of FliP sequences from *Salmonella* and various species of the *Vibrionales* order. Red indicates the signal peptide, yellow the N-terminal domain and light blue the part of the sequence that was too poorly ordered to model in the structures of *S.* Typhimurium and *V. mimicus* FliPQR.

**Supplementary Fig. 2 Data processing strategy for *V. mimicus* FliPQR**

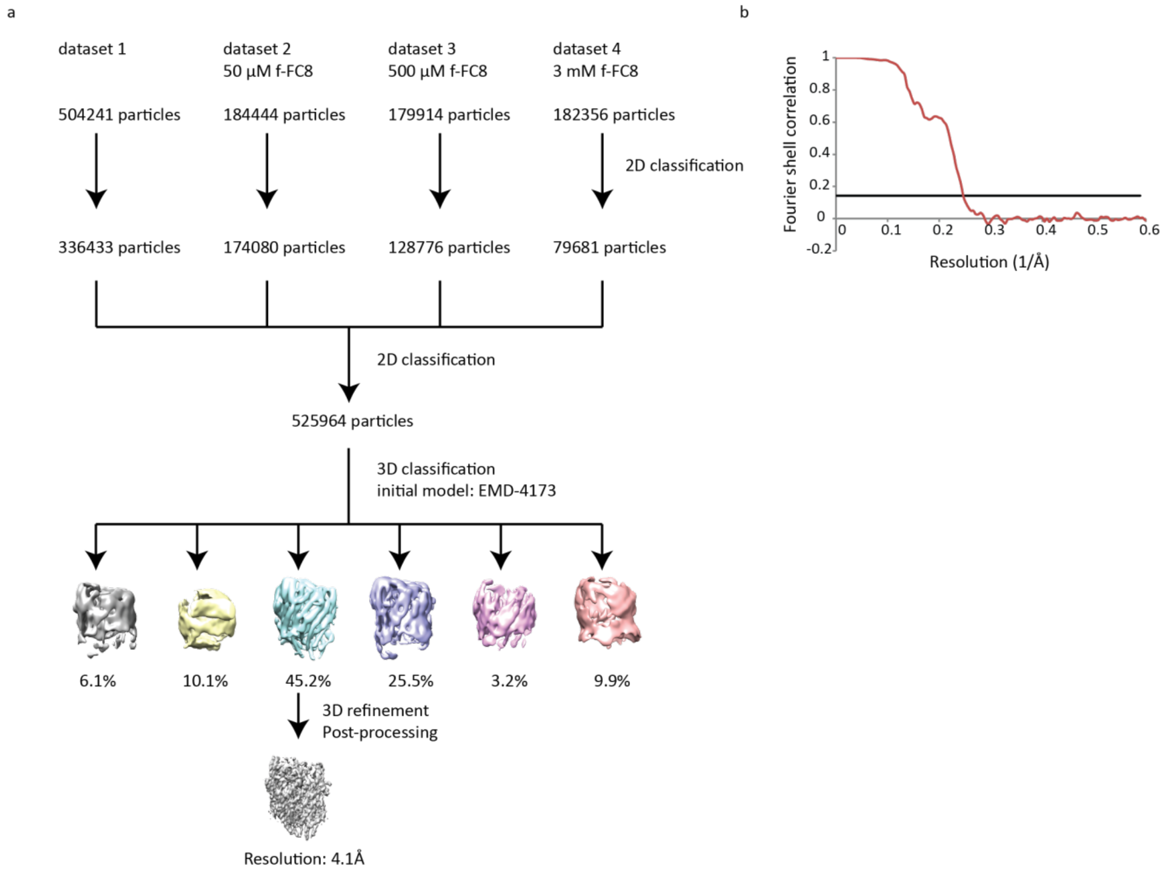

**a**, Single particle averaging of cryo-EM data of *V. mimicus* FliPQR. **b**, Gold standard FSC curve.

**Supplementary Fig. 3 Data processing strategy for *P. savastanoi* FliPQR**

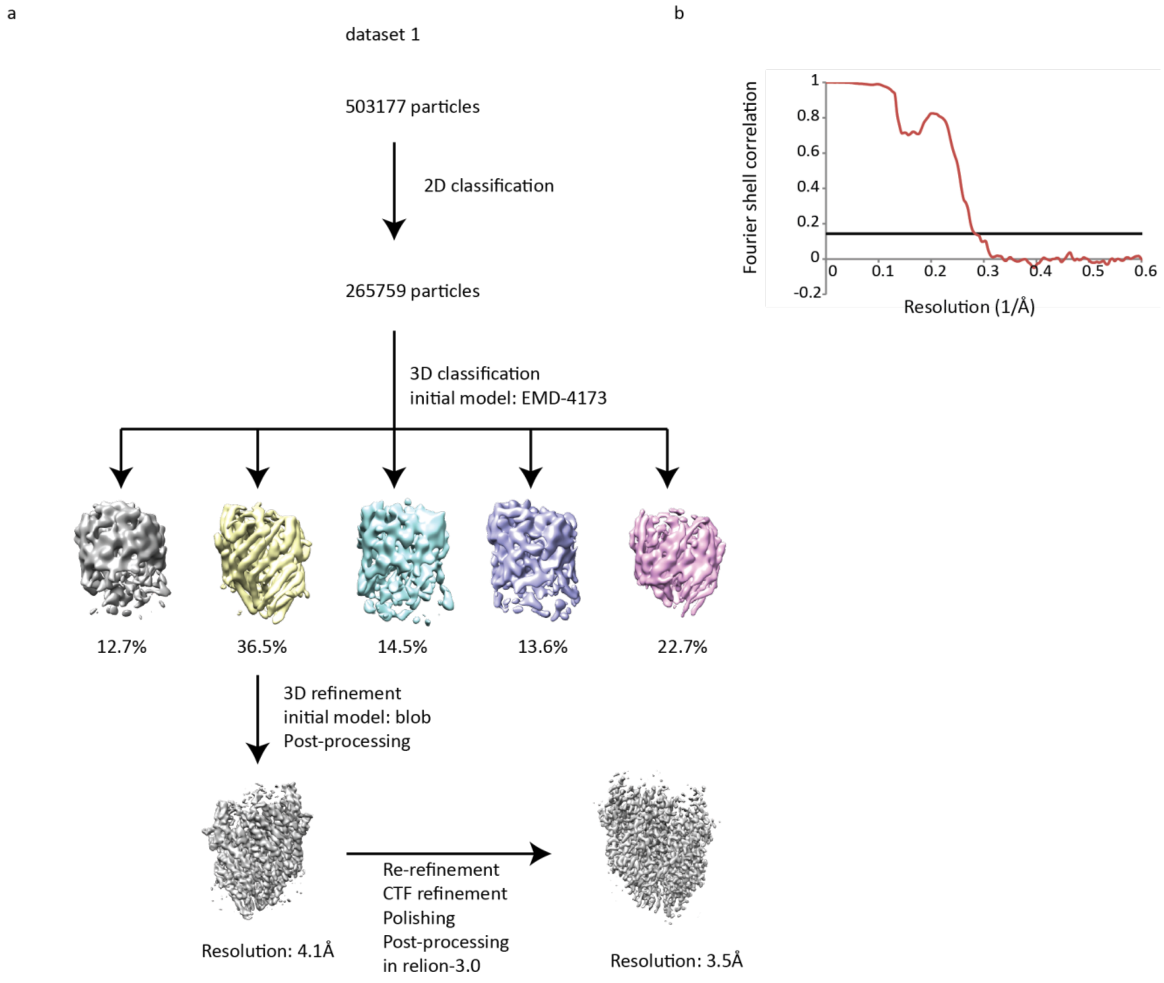

**a**, Single particle averaging of cryo-EM data of *P. savastanoi* FliPQR. **b**, Gold standard FSC curve.

**Supplementary Fig. 4 Mapping the interaction site of SctU by *in vivo* photocrosslinking**

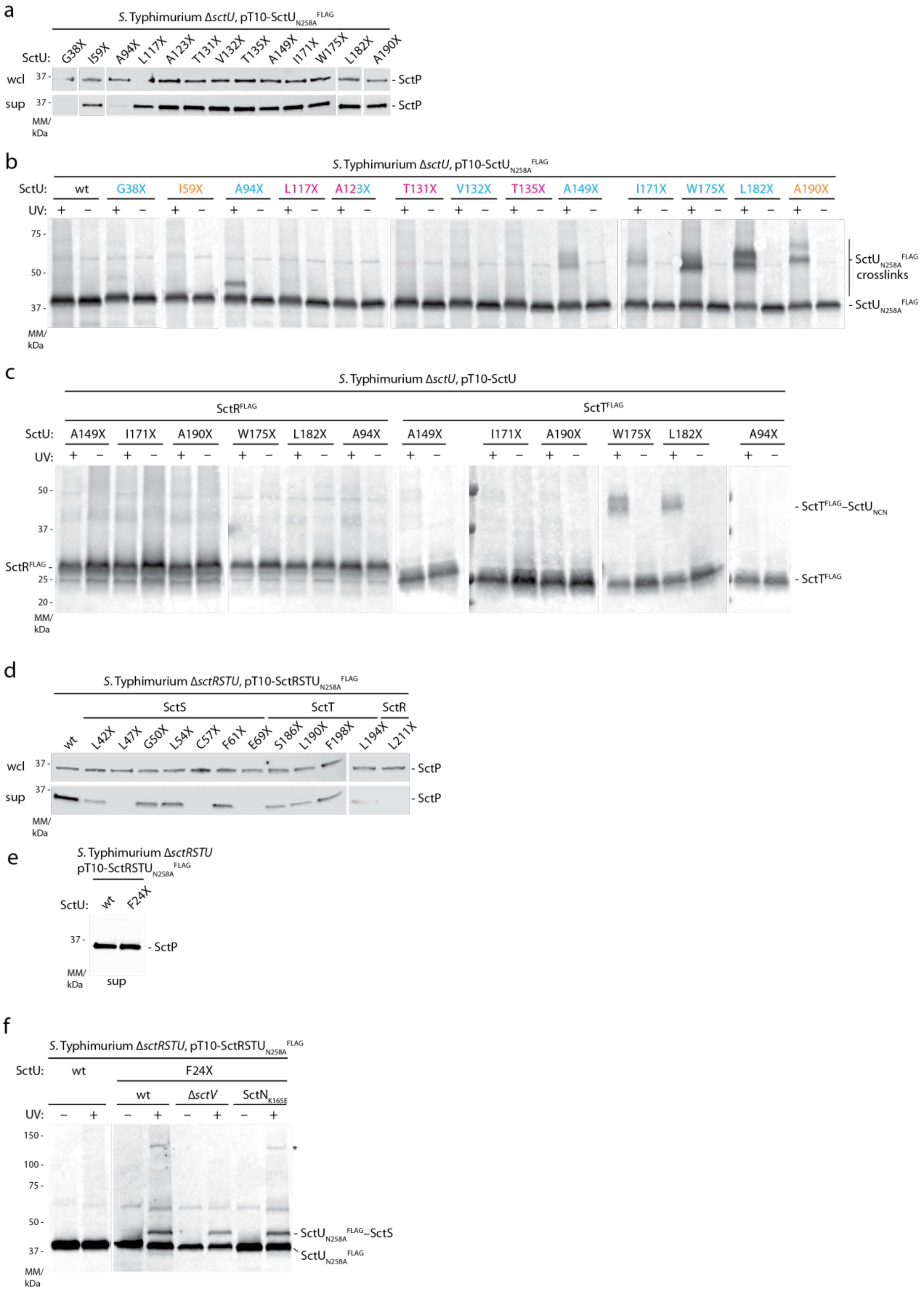

Tested were interaction sites between SctU and SctR, SctS, and SctT, respectively, that where predicted by covariance analysis. **a**, Immunodetection of the early T3SS substrate SctP on Western blots of SDS PAGE-separated whole cell lysates and culture supernatants of the indicated S. Typhimurium SctU pBpa mutants. **b**, Immunodetection of SctU$^{FLAG}$ on Western blots of SDS PAGE-separated crude membrane samples of the indicated S. Typhimurium SctU pBpa mutants (denoted with X). Each sample is shown with and without UV-irradiation to induce photocrosslinking of pBpa to neighboring interaction partners. Colors indicate covariance predition of SctU to SctR (orange), SctS (magenta), or SctT (cyan). **c**, As in b but showing immunodetection of SctR$^{FLAG}$ or SctT$^{FLAG}$. **d**, As in a but assessing secretion of SctR, SctS, and SctT pBpa mutants, respectively. **e**, Immunodetection of the early T3SS substrate SctP on Western blots of SDS PAGE-separated culture supernatants of the S. enterica wild type and a SctU F24pBpa (denoted as X) mutant. **f**, Immunodetection of SctU$^{FLAG}$ on Western blots of SDS PAGE-separated crude membrane samples of S. enterica wild type and a SctU F24pBpa mutant. The SctU F24X crosslink was assessed in the wild type, in a ΔsctV and in an ATP hydrolysis-deficient SctNK165E mutant strain. The asterisk indicates the presence of a putative SctU-SctV crosslink.

**Supplementary Fig. 5 Data processing strategy for *V. mimicus* FliPQR-FlhB**

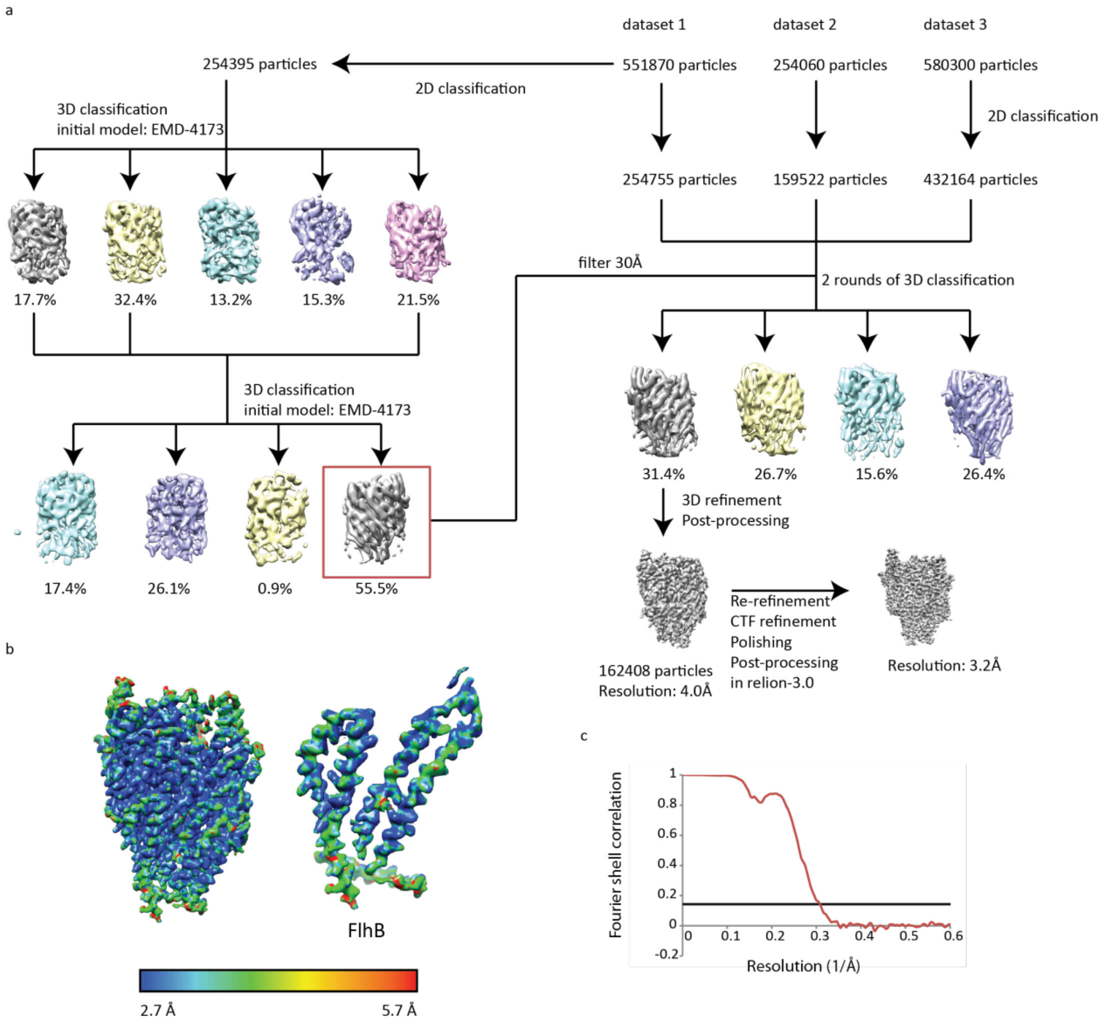

**a**, Single particle averaging of cryo-EM data of *V. mimicus* FliPQR-FlhB. **b**, Surface representation of the unsharpened volume coloured by local resolution calculated in ResMap (*34*). Entire map (left) and zoom of the volume corresponding to FlhB (right). **c**, Gold standard FSC curve.

**Supplementary Fig. 6 Potential lipidic ligands.**

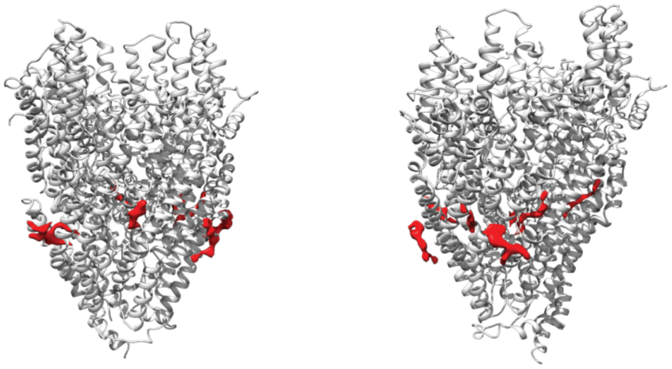

Structure of FliPQR (white) and unmodelled density (red).

**Supplementary Fig. 7 Co-variation within FlhB**

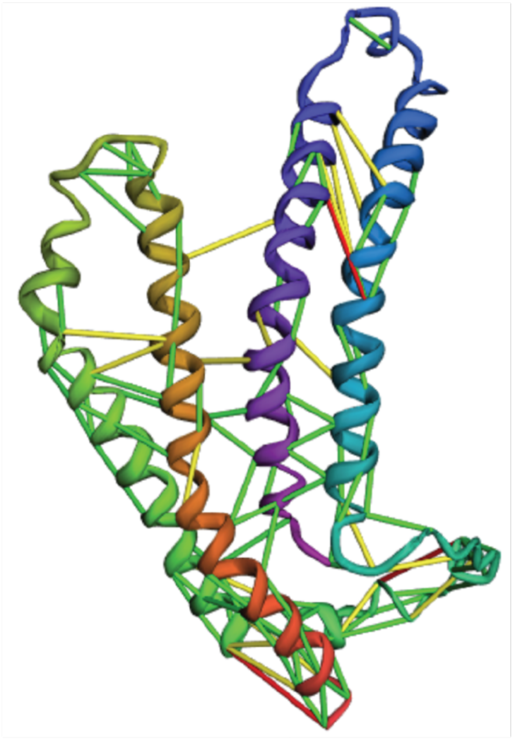

FlhB model with lines connecting co-varying residues. Co-variation was calculated in gremlin (*35*) and only contacts with a gremlin score higher than 0.75 are displayed. Green lines indicate co-varying residues with a distance < 5 Å, yellow lines 5 – 10 Å and red > 10 Å.

**Supplementary Fig. 8 Conservation within the FliPQR-FlhB complex**

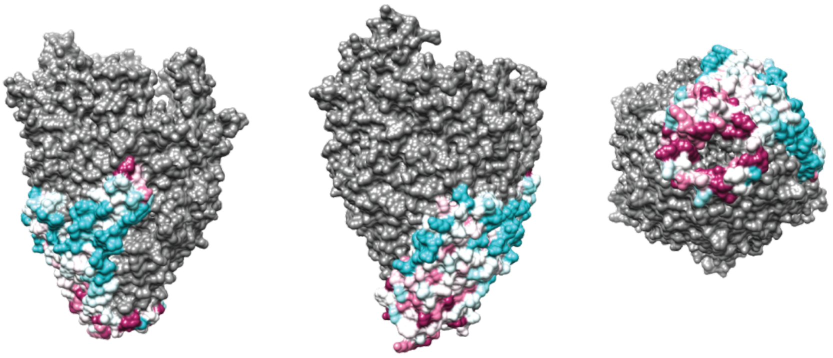

ConSurf (*36*) representation of sequence conservation of FlhB shows that FlhB$_{L}$ is highly conserved. Magenta represents highly conserved residues and light blue highly variable residues.

**Supplementary Fig. 9 The poorly resolved FlhB N-terminus and FlhB$_{CN}$ are directly underneath the entrance to the lumen of the FliPQR-FlhB complex**

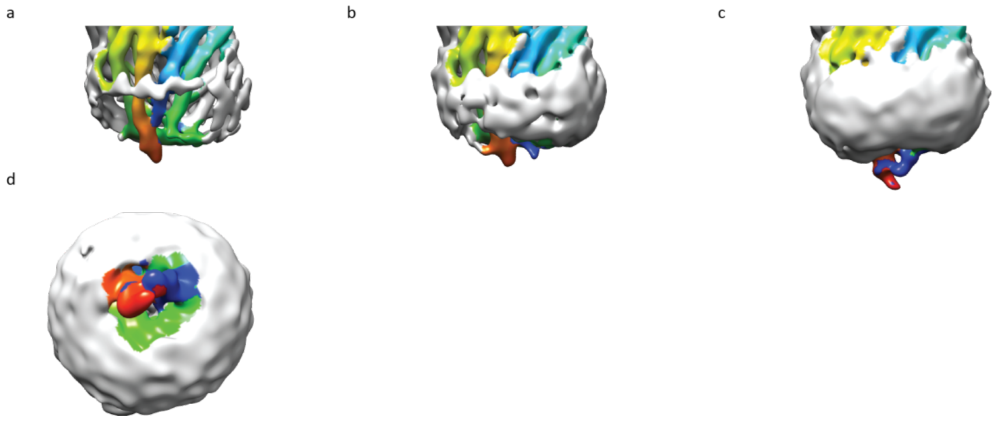

**a,b,c**, The FliPQR-FlhB cryo-EM density map, filtered to 8 Å, is shown at three different contour levels. Density corresponding to FlhB is coloured in rainbow colours. **d**, View from the cytoplasm shows that the termini of FlhB$_{TM}$ block access to the gate.

**Supplementary Fig. 10 Motility of *E. coli* W strains**

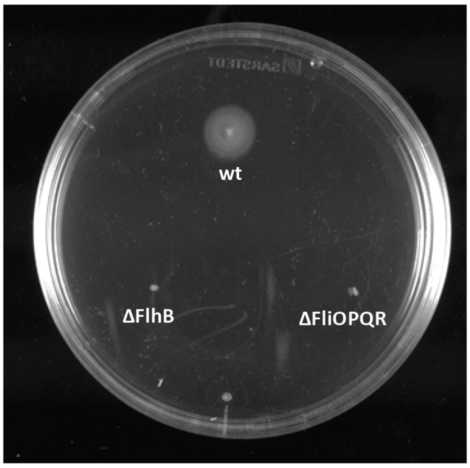

Motility of *E. coli* W wild type, strain WL1 (ΔFliOPQR, right) and strain WL2 (ΔFlhB, left) in soft agar.

**Supplementary Fig. 11 Purification of mutant FliPQR FlhB complexes**

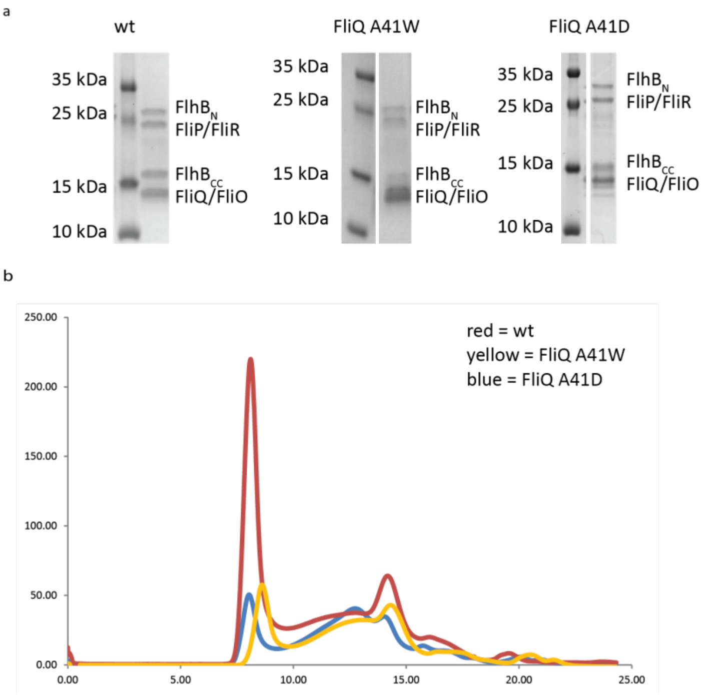

**a**, SDS-PAGE analysis of the StrepTrap eluate of FliOPQR-FlhB wild type (left, same as **Error! Reference source not found.**a), FliQ A41W mutant (middle) and FliQ A41D mutant (right). **b**, Analytical SEC of wild type (red curve), FliQ A41W (yellow curve) and FliQ A41D (blue curve) complexes.

**Supplementary Fig. 12 Photocrosslinking of SctU**

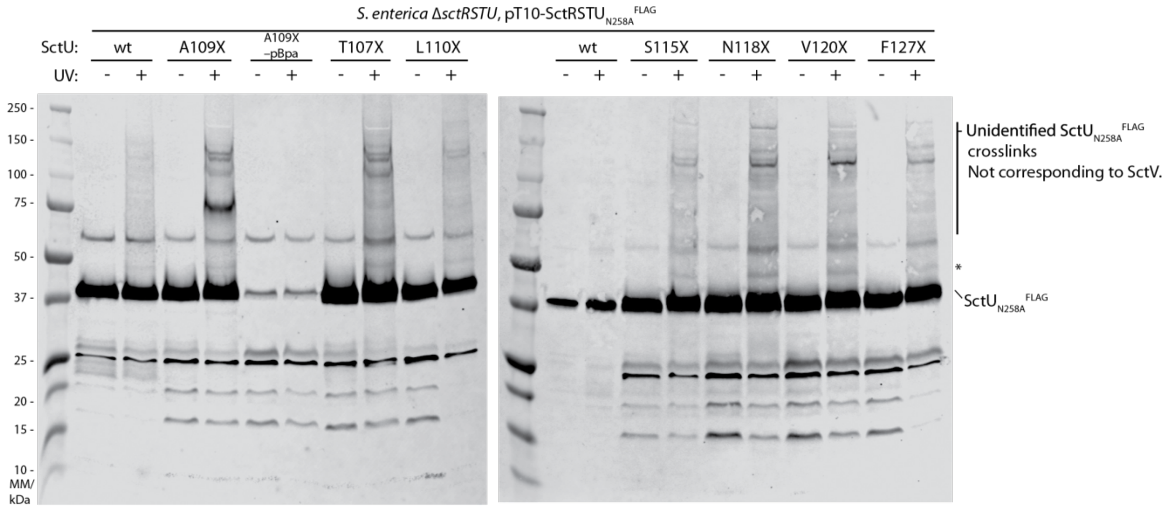

Immunodetection of SctU$^{FLAG}$ on Western blots of SDS PAGE-separated crude membrane samples of the indicated *S*. Typhimurium SctU *p*Bpa mutants (denoted with X). Each sample is shown with and without UV-irradiation to induce photo crosslinking of *p*Bpa to neighboring interaction partners. The – *p*Bpa control shows that little skipping of the amber stop codon occurs in the absence of *p*Bpa. The crosslinks that were specific to the *p*Bpa mutants could not be assigned to SctV or SctS. The asterisk shows a putative SctU-SctS crosslink, which, however, is also slightly visible in one wild type control.

Supplementary tables

**Supplementary table 1**

| SctU *S.* Typhimurium | FlhB *V. mimicus* | FlhB *E. coli* W | | SctS *S.* Typhimurium | FliQ *V. mimicus* | FliQ *E. coli* W |
|---|---|---|---|---|---|---|
| F24 | A28 | P27 | | T38 | A40 | A40 |
| K27 | K31 | R30 | | V39 | A41 | A41 |
| D28 | E32 | E31 | | T40 | T42 | T42 |
| G38 | G42 | G41 | | Q41 | S43 | Q43 |
| I59 | Q62 | S61 | | L42 | I44 | I44 |
| A94 | L105 | L104 | | L47 | L49 | L49 |
| T107 | A118 | G117 | | G50 | L52 | I52 |
| A109 | A120 | S119 | | L54 | I56 | I56 |
| L110 | A121 | L120 | | C57 | L59 | F59 |
| S115 | S126 | S125 | | F61 | M63 | I63 |
| L117 | M128 | L127 | | E69 | Q71 | N71 |
| N118 | N129 | N128 | | | | |
| V120 | L131 | L130 | | **SctT *S.* Typhimurium** | **FliR *V. mimicus*** | **FliR *E. coli* W** |
| G122 | G133 | G132 | | S186 | L189 | L188 |
| A123 | F134 | I133 | | L190 | T193 | I192 |
| K125 | R136 | R135 | | L194 | T197 | T196 |
| L126 | M137 | M136 | | F198 | S201 | A200 |
| F127 | F138 | F137 | | | | |
| T131 | S142 | T141 | | **SctR *S.* Typhimurium** | **FliP *V. mimicus*** | **FliP *E. coli* W** |
| V132 | W143 | G142 | | L211 | L288 | L234 |
| D134 | E145 | E144 | | | | |
| T135 | L146 | L145 | | | | |
| K137 | K148 | K147 | | | | |
| S143 | K152 | K151 | | | | |
| A149 | G158 | S157 | | | | |
| I171 | H185 | N184 | | | | |
| W175 | L187 | M186 | | | | |
| L182 | F193 | C192 | | | | |
| A190 | L201 | V200 | | | | |
| D197 | D208 | D207 | | | | |

**Supplementary table 2**

| name | number | sequence |
|---|---|---|
| pT12_f | 1 | GAAAACCTGTACTTCCAGGGTC |
| pT12_r | 2 | TGGTGAATTCCTCCTGAATTTC |
| FliO_VIBMI_f | 3 | TCAGGAGGAATTCACCATGCGAAAATTGGCGGCTCTG |
| FliR_VIBMI_r | 4 | GGAAGTACAGGTTTTCACAGTTCAAGCGAATGAGACGACAGATCTGCTC |
| FlhB_VIBMI_r | 5 | GGAAGTACAGGTTTTCGTAACGCATATCTGGTGGGATCGGCATATTCTC |
| FliO_ECOLW_f | 6 | TTCAGGAGGAATTCACCATGAATAACCACGCTACTGTG |
| FliR_ECOLW_r | 7 | TGGAAGTACAGGTTTTCTATTAATGGCAATTCACTAATAATATCAGC |
| FlhB_ ECOLW_1_f | 8 | TTCAGGAGGAATTCACCGTGTCTGACGAGAGCGACGACAAAACAG |
| FlhB_ECOLW_1_r | 9 | TGGAAGTACAGGTTTTCCTCATGGGTCGGTTTCTCG |
| FlhB_ECOLW_2_f | 10 | AACAGGAGGAATTAACCGTGTCTGACGAGAGCGACGACAAAACAG |
| FlhB_ECOLW2_r | 11 | GATGGTCGACGGCGCTTTATTTTTCGAACTGCGGGTGGCTCCACGATCCACCTCCCGATCCACCTCCGGAAC |
| FliQ ECOLW A41W f | 12 | TGCAGGCCTGGACGCAGATCAACGAAATGACGCTGTCGTTTATTCC |
| FliQ ECOLW A41W r | 13 | ATCTGCGTCCAGGCCTGCAAAATACTGATGATAAGGCCCGTGACCAGCGCCACCAGCAAC |
| FliQ ECOLW A41D f | 14 | TGCAGGCCGACACGCAGATCAACGAAATGACGCTGTCGTTTATTCC |
| FliQ ECOLW A41D r | 15 | ATCTGCGTGTCGGCCTGCAAAATACTGATGATAAGGCCCGTGACCAGCGCCACCAGCAAC |
| FliQ ECOLW A41R f | 16 | TGCAGGCCCGCACGCAGATCAACGAAATGACGCTGTCGTTTATTCC |
| FliQ ECOLW A41R r | 17 | ATCTGCGTGCGGGCCTGCAAAATACTGATGATAAGGCCCGTGACCAGCGCCACCAGCAAC |
| FliQ ECOLW delta A41 f | 18 | TGCAGGCCACGCAGATCAACGAAATGACGCTGTCGTTTATTCC |
| FliQ ECOLW delta A41 r | 19 | ATCTGCGTGGCCTGCAAAATACTGATGATAAGGCCCGTGACCAGCGCCACCAGCAAC |
| FliQ ECOLW T42W f | 20 | TGCAGGCCGCCTGGCAGATCAACGAAATGACGCTGTCGTTTATTCC |
| FliQ | 21 | ATCTGCCAGGCGGCCTGCAAAATACTGATGATAAGGCCCGTGACCAGCGCCACCAGCAAC |

| | | |
|---|---|---|
| ECOLW T42W r | | |
| FliQ ECOLW T42D f | 22 | TGCAGGCCGCCGACCAGATCAACGAAATGACGCTGTCGTTTATTCC |
| FliQ ECOLW T42D r | 23 | ATCTGGTCGGCGGCCTGCAAAATACTGATGATAAGGCCCGTGACCAGCGCCACCAGCAAC |
| FlhB ECOLW L113 deletion r | 24 | GCCAGAACCGCCTCCCCCTCCCAGCATGACCGGAGAAATGAGCGCCACCAGCAC |
| FlhB ECOLW S138 deletion f | 25 | GGAGGCGGTTCTGGCTCGGCTCAGACTGGCGCGGAGTTGCTTAAAGC |
| pBAD_f | 26 | AGCGCCGTCGACCATCATC |
| pBAD_r | 27 | CATGGTTAATTCCTCCTGTTAGCCC |
| FlhB ECOLW G123A f | 28 | CGCTACCGGCCATTAAACGGATGTTCTCGGCTCAGACTG |
| FlhB ECOLW G123A r | 29 | CGTTTAATGGCCGGTAGCGGGTTGAGTTTGGAAAACTTCG |
| FlhB ECOLW G123D f | 30 | CGCTACCGGACATTAAACGGATGTTCTCGGCTCAGACTG |
| FlhB ECOLW G123D r | 31 | CGTTTAATGTCCGGTAGCGGGTTGAGTTTGGAAAACTTCG |
| FlhB ECOLW I133 deletion f | 32 | TACCGGGCATGTTCTCGGCTCAGACTGGCGCGGAGTTGCTTAAAGC |
| FlhB ECOLW R135 deletion r | 33 | GAGAACATGCCCGGTAGCGGGTTGAGTTTGGAAAACTTCG |
| FlhB ECOLW Q121 deletion1 f | 34 | ATTTAGCGGCCAGCCGAAGTTTTCCAAACTCAACCCGCTACCGGGCATTAAACG |
| FlhB ECOLW G117 deletion r | 35 | ACTTCGGCTGGCCGCTAAATACCAGCCCTCCCAGCATGACCGGAGAAATGAG |
| FlhB ECOLW Q121 deletion2 f | 36 | AGGGCTGGTACAGCCGAAGTTTTCCAAACTCAACCCGCTACCGGGCATTAAACG |
| FlhB ECOLW V114 | 37 | ACTTCGGCTGTACCAGCCCTCCCAGCATGACCGGAGAAATGAGC |

| | | |
|---|---|---|
| deletion r | | |
| FliOPQR KO f | 38 | CACCGATATCATTACTCCGTCTGAGCGAATGCGTCGCCTGAGCCGTTAGTGATGAATAACGAAGTTCCTATACTTTCTAGAGAATAGGAACTTCggaataggaacttc |
| FliOPQR KO r | 39 | TCGCAGGTATTATTTTTCGGATAATCCTTAGGATAACATGATAAACGTTACGGAATTATAGAAGTTCCTATTCTCTAGAAAGTATAGGAACTTCggcgcgcctacctgtg |
| FlhB KO f1 | 40 | TGTCTGCGAAGTGGTCGATCTTAGCCAGGTAAGC |
| FlhB KO r1 | 41 | CTGGAAAGAGTATTAAGCGTGAATGATGCCAGAGCGCAAAGC |
| FlhB KO f2 | 42 | TCATTCACGCTTAATACTCTTTCCAGGATTGGCGACGTGTCTGACGAGAGGAAGTTCCTATACTTTCTAGAGAATAGGAACTTCggaataggaacttc |
| FlhB KO r2 | 43 | GCATCGCGGCCAGATTACTCATGGGTCGGTTTCTCGTTAATAAAATCCAGGAAGTTCCTATTCTCTAGAAAGTATAGGAACTTCggcgcgcctacctgtg |
| FlhB KO f3 | 44 | GACCCATGAGTAATCTGGCCGCGATGCTG |
| FlhB KO r3 | 45 | ACTTTCAATCCAGATAGCATTCAGGCCAAATGC |
| SpaQ_T38X_QC_f | 46 | CTGGTAGGGTTATTCCAGTAGGTAACGCAATTACAGGAACAG |
| SpaQ_T38X_QC_r | 47 | CTGTTCCTGTAATTGCGTTACCTACTGGAATAACCCTACCAG |
| SpaQ_V39X_QC_f | 48 | GGTAGGGTTATTCCAGACGTAGACGCAATTACAGGAACAGACG |
| SpaQ_V39X_QC_r | 49 | CGTCTGTTCCTGTAATTGCGTCTACGTCTGGAATAACCCTACC |
| SpaQ_T40X_QC_f | 50 | GGGTTATTCCAGACGGTATAGCAATTACAGGAACAGACG |
| SpaQ_T40X_QC_r | 51 | CGTCTGTTCCTGTAATTGCTATACCGTCTGGAATAACCC |
| SpaQ_Q41X_QC_f | 52 | GTTATTCCAGACGGTAACGTAGTTACAGGAACAGACGCTG |
| SpaQ_Q41X_QC_r | 53 | CAGCGTCTGTTCCTGTAACTACGTTACCGTCTGGAATAAC |
| QC_SpaQ_L42X_f | 54 | CCAGACGGTAACGCAATAGCAGGAACAGACGCTG |
| QC_SpaQ_L42X_r | 55 | CAGCGTCTGTTCCTGCTATTGCGTTACCGTCTGG |
| SpaS_A94X_QC_f | 56 | CTGCTCTGCTTAGTGTGTTCCTAGTTACCGGCGTTATTACAGGC |
| SpaS_A94X_QC_r | 57 | GCCTGTAATAACGCCGGTAACTAGGAACACACTAAGCAGAGCAG |
| SpaS_T107X_QC_f | 58 | CGGTTTTGTGCTGGCGTAGGAAGCATTAAAGCCTAATTTATCGGC |
| SpaS_T107X_QC_r | 59 | GCCGATAAATTAGGCTTTAATGCTTCCTACGCCAGCACAAAACCG |
| SpaS_A109X_QC_f | 60 | GTTTTGTGCTGGCGACAGAATAGTTAAAGCCTAATTTATCGGCGTTAAACC |
| SpaS_A109X_QC_r | 61 | GGTTTAACGCCGATAAATTAGGCTTTAACTATTCTGTCGCCAGCACAAAAC |
| SpaS_L110X_QC_f | 62 | CTGGCGACAGAAGCATAGAAGCCTAATTTATCGGCGTTAAAC |
| SpaS_L110X_QC_r | 63 | GTTTAACGCCGATAAATTAGGCTTCTATGCTTCTGTCGCCAG |
| SpaS_S115 | 64 | CAGAAGCATTAAAGCCTAATTTATAGGCGTTAAACCCGGTAG |

| | | |
|---|---|---|
| X_QC_f | | |
| SpaS_S115 X_QC_r | 65 | CTACCGGGTTTAACGCCTATAAATTAGGCTTTAATGCTTCTG |
| SpaS_N11 8X_QC_f | 66 | GCCTAATTTATCGGCGTTATAGCCGGTAGAAGGGGCAAAAAAAC |
| SpaS_N11 8X_QC_r | 67 | GTTTTTTTGCCCCTTCTACCGGCTATAACGCCGATAAATTAGGC |
| SpaS_V120 X_QC_f | 68 | CTAATTTATCGGCGTTAAACCCGTAGGAAGGGGCAAAAAAACTTTTTAGTATGC |
| SpaS_V120 X_QC_r | 69 | GCATACTAAAAAGTTTTTTTGCCCCTTCCTACGGGTTTAACGCCGATAAATTAG |
| SpaS_F127 X_QC_f | 70 | GGTAGAAGGGGCAAAAAAACTTTAGAGTATGCGCACGGTTAAAG |
| SpaS_F127 X_QC_r | 71 | CTTTAACCGTGCGCATACTCTAAAGTTTTTTTGCCCCTTCTACC |
| QC_SpaQ_ L47X_f | 96 | GCAATTACAGGAACAGACGTAGCCTTTTGGCATTAAATTACTTGG |
| QC_SpaQ_ L47X_r | 97 | CCAAGTAATTTAATGCCAAAAGGCTACGTCTGTTCCTGTAATTGC |
| QC_SpaQ_ G50X_f | 98 | GGAACAGACGCTGCCTTTTTAGATTAAATTACTTGGCGTGTGTTTATGC |
| QC_SpaQ_ G50X_r | 99 | GCATAAACACACGCCAAGTAATTTAATCTAAAAAGGCAGCGTCTGTTCC |
| QC_SpaQ_ L54X_f | 100 | GACGCTGCCTTTTGGCATTAAATTATAGGGCGTGTGTTTATGCTTG |
| QC_SpaQ_ L54X_r | 101 | CAAGCATAAACACACGCCCTATAATTTAATGCCAAAAGGCAGCGTC |
| QC_SpaQ_ C57X_f | 102 | GGCATTAAATTACTTGGCGTGTAGTTATGCTTGTTTTTACTGTCTGG |
| QC_SpaQ_ C57X_r | 103 | CCAGACAGTAAAAACAAGCATAACTACACGCCAAGTAATTTAATGCC |
| QC_SpaQ_ F61X_f | 104 | GGCGTGTGTTTATGCTTGTAGTTACTGTCTGGCTGGTATG |
| QC_SpaQ_ F61X_r | 105 | CATACCAGCCAGACAGTAACTACAAGCATAAACACACGCC |
| QC_SpaQ_ E69X_f | 106 | GTCTGGCTGGTATGGCTAGGTTTTACTCTCTTACGGG |
| QC_SpaQ_ E69X_r | 107 | CCCGTAAGAGAGTAAAACCTAGCCATACCAGCCAGAC |
| QC_SpaR_ S186X_f | 108 | GCCTTGGTTCTGGCCTAGCCGGTGGTATTAGTGCTG |
| QC_SpaR_ S186X_r | 109 | CAGCACTAATACCACCGGCTAGGCCAGAACCAAGGC |
| QC_SpaR_ L190X_f | 110 | GCCAGTCCGGTGGTATAGGTGCTGTTGCTGTCAG |
| QC_SpaR_ L190X_r | 111 | CTGACAGCAACAGCACCTATACCACCGGACTGGC |
| QC_SpaR_ F198X_f | 112 | GCTGTTGCTGTCAGAAGTATAGCTGGGTTTATTGTCGCG |
| QC_SpaR_ F198X_r | 113 | CGCGACAATAAACCCAGCTATACTTCTGACAGCAACAGC |

| | | |
|---|---|---|
| QC_SpaR_ L194X_f | 114 | GGTGGTATTAGTGCTGTTGTAGTCAGAAGTATTCCTGGG |
| QC_SpaR_ L194X_r | 115 | CCCAGGAATACTTCTGACTACAACAGCACTAATACCACC |
| QC_SpaP_ L211X_f | 116 | CTTGATGGCTGGACCTTATAGTCTAAGGGATTGATATTACAGTATATG |
| QC_SpaP_ L211X_r | 117 | CATATACTGTAATATCAATCCCTTAGACTATAAGGTCCAGCCATCAAG |
| SpaS_QC_ F24X_f | 118 | CCGCTAAAAAAGGCCAGTCATAGAAAAGTAAAGATCTCATTATCGC |
| SpaS_QC_ F24X_r | 119 | GCGATAATGAGATCTTTACTTTTCTATGACTGGCCTTTTTTAGCGG |
| FlhB ECOLW R135A f | 120 | ATTAAAGCGATGTTCTCGGCTCAGACTGGCGCGGAGTTG |
| FlhB ECOLW R135A r | 121 | GAGCCGAGAACATCGCTTTAATGCCCGGTAGCGGGTTGAG |
| FlhB ECOLW L127A f | 122 | GTTTTCCAAAGCCAACCCGCTACCGGGCATTAAACGGATGTTCTC |
| FlhB ECOLW L127A r | 123 | TTAATGCCCGGTAGCGGGTTGGCTTTGGAAAACTTCGGCTGCAAGGATTTGC |
| FlhB ECOLW L127D f | 124 | GTTTTCCAAAGACAACCCGCTACCGGGCATTAAACGGATGTTCTC |
| FlhB ECOLW L127D r | 125 | TTAATGCCCGGTAGCGGGTTGTCTTTGGAAAACTTCGGCTGCAAGGATTTGC |
| FlhB ECOLW M136A f | 126 | CGCTACCGGGCATTAAACGGGCGTTCTCGGCTCAGACTGGCGCGGAGTTG |
| FlhB ECOLW M136D f | 127 | CGCTACCGGGCATTAAACGGGACTTCTCGGCTCAGACTGGCGCGGAGTTG |

**Supplementary table 3**

| strains | | | |
|---|---|---|---|
| name | description | reference | construction note |
| *E. coli W* | DSM 1116 | | |
| WL1 | *E. coli W* ΔFliOPQR | this study | Made by λ Red recombination of the wild type strain with the PCR product of the following primers and template: Primers 38 and 39 and pKD3 |
| WL2 | *E. coli W* ΔFlhB | this study | Made by λ Red recombination of the wild type strain with the Gibson assembly product of the PCR products of the following primer/template pairs: Primers 42 and 43 and pKD3, primers 40 and 41 with *E. coli* W genomic DNA and primers 44 and 45 with *E. coli* W genomic DNA. |
| plasmids | | | |
| name | description | reference | construction note |
| pMIB5689 | pT12, SpaPQR$^{TEV2xStrepII}$ | Dietsche, T., Tesfazgi Mebrhatu, M., Brunner, M. J., Abrusci, P., Yan, J., Franz-Wachtel, M., et al. (2016). Structural and functional characterization of the bacterial type III secretion export apparatus. PLoS Pathogens, 12(12), e1006071. http://doi.org/10.1371/journal.ppat.1006071 | |
| pT12_FliOPQR_PSESH | pT12, *P. savastanoi* FliOPQR$^{TEV2xStrepII}$ | Kuhlen L., Abrusci P., Johnson S., Gault J., Deme J., Caesar J., Dietsche T., Mebrhatu M. T., Ganief T., Macek B., Wagner S., Robinson C. V., Lea S. M. (2018). Structure of the core of the type III secretion system export apparatus. Nat Struct Mol Biol. 25(7):583-590. https://doi.org/10.1038/s41594-018-0086-9 | |
| pT12_FliOPQR_VIBMI | pT12, *V. mimicus* FliOPQR$^{TEV2xStrepII}$ | this study | Made by Gibson assembly of PCR products of the following two |

| | | | |
|---|---|---|---|
| | | | primer/template pairs: 1. Primers 1 and 2 from pMIB5689; 2. primers 3 and 4 from *V. mimicus* genomic DNA. |
| pT12_FliOPQR_FlhB_VIBMI | pT12, *V. mimicus* FliOPQR $FlhB^{TEV2xStrepII}$ | this study | Made by Gibson assembly of PCR products of the following two primer/template pairs: 1. Primers 1 and 2 from pMIB5689; 2. primers 3 and 5 from *V. mimicus* genomic DNA. |
| pT12_W1 | pT12, *E. coli* W $FliOPQR^{TEV2xStrepII}$ | this study | Made by *in vivo* assembly of PCR products of the following two primer/template pairs: 1. Primers 1 and 2 from pMIB5689; 2. primers 6 and 7 from E. *coli* W genomic DNA. |
| pT12_W2 | pT12, *E. coli* W $FliOPQR^{TEV2xStrepII}$ FliQ A41W | this study | Made by *in vivo* assembly of the PCR product of the following primer/template pair: Primers 12 and 13 from pT12_W1. |
| pT12_W3 | pT12, *E. coli* W $FliOPQR^{TEV2xStrepII}$ FliQ A41D | this study | Made by *in vivo* assembly of the PCR product of the following primer/template pair: Primers 14 and 15 from pT12_W1. |
| pT12_W4 | pT12, *E. coli* W $FliOPQR^{TEV2xStrepII}$ FliQ A41R | this study | Made by *in vivo* assembly of the PCR product of the following primer/template pair: Primers 16 and 17 from pT12_W1. |
| pT12_W5 | pT12, *E. coli* W $FliOPQR^{TEV2xStrepII}$ FliQ ΔA41 | this study | Made by *in vivo* assembly of the PCR product of the following primer/template pair: Primers 18 and 19 from pT12_W1. |
| pT12_W6 | pT12, *E. coli* W $FliOPQR^{TEV2xStrepII}$ FliQ T42W | this study | Made by *in vivo* assembly of the PCR product of the following primer/template pair: Primers 20 and 21 from pT12_W1. |
| pT12_W7 | pT12, *E. coli* W | this study | Made by *in vivo* |

| | | | |
|---|---|---|---|
| | FliOPQR$^{TEV2xStrepII}$ FliQ T42D | | assembly of the PCR product of the following primer/template pair: Primers 22 and 23 from pT12_W1. |
| pT12_W8 | pT12, *E. coli* W FlhB$^{TEV2xStrepII}$ | this study | Made by *in vivo* assembly of PCR products of the following two primer/template pairs: 1. Primers 1 and 2 from pMIB5689; 2. primers 8 and 9 from E. *coli* W genomic DNA. |
| pBAD_W1 | pBAD, *E. coli* W FlhB$^{TEV2xStrepII}$ | this study | Made by *in vivo* assembly of PCR products of the following two primer/template pairs: 1. Primers 26 and 27 from pSZ2; 2. primers 10 and 11 from pT12_W8. |
| pBAD_W2 | pBAD, *E. coli* W FlhB$^{TEV2xStrepII}$ G132A | this study | Made by *in vivo* assembly of the PCR product of the following primer/template pair: Primers 28 and 29 from pBAD_W1. |
| pBAD_W3 | pBAD, *E. coli* W FlhB$^{TEV2xStrepII}$ G132D | this study | Made by *in vivo* assembly of the PCR product of the following primer/template pair: Primers 30 and 31 from pBAD_W1. |
| pBAD_W4 | pBAD, *E. coli* W FlhB$^{TEV2xStrepII}$ Δ133-135 | this study | Made by *in vivo* assembly of the PCR product of the following primer/template pair: Primers 32 and 33 from pBAD_W1. |
| pBAD_W5 | pBAD, *E. coli* W FlhB$^{TEV2xStrepII}$ Δ118-120 | this study | Made by *in vivo* assembly of the PCR product of the following primer/template pair: Primers 34 and 35 from pBAD_W1. |
| pBAD_W6 | pBAD, *E. coli* W FlhB$^{TEV2xStrepII}$ Δ115-120 | this study | Made by *in vivo* assembly of the PCR product of the following primer/template pair: Primers 36 and 37 from pBAD_W1. |
| pBAD_W7 | pBAD, *E. coli* W FlhB$^{TEV2xStrepII}$ | this study | Made by *in vivo* assembly of the PCR |

| | | | |
|---|---|---|---|
| | Δ113-137 | | product of the following primer/template pair: Primers 24 and 25 from pBAD_W1. |
| pBAD_W8 | pBAD, E. coli W FlhBTEV2xStrep II R135A | this study | Made by in vivo assembly of the PCR product of the following primer/template pair: Primers 120 and 121 from pBAD_W1. |
| pBAD_W9 | pBAD, E. coli W FlhBTEV2xStrep II L127A | this study | Made by in vivo assembly of the PCR product of the following primer/template pair: Primers 122 and 123 from pBAD_W1. |
| pBAD_W10 | pBAD, E. coli W FlhBTEV2xStrep II L127D | this study | Made by in vivo assembly of the PCR product of the following primer/template pair: Primers 124 and 125 from pBAD_W1. |
| pBAD_W11 | pBAD, E. coli W FlhBTEV2xStrep II L127A M136A | this study | Made by in vivo assembly of the PCR product of the following primer/template pair: Primers 123 and 126 from pBAD_W1. |
| pBAD_W12 | pBAD, E. coli W FlhBTEV2xStrep II L127D M136D | this study | Made by in vivo assembly of the PCR product of the following primer/template pair: Primers 125 and 127 from pBAD_W1. |
| pSZ2 | pBAD, *S. flexneri* MxiG$^{myc\text{-}His}$ | Zenk S. F., Stabat D., Hodgkinson J. L., Veenendaal A. K., Johnson S., Blocker A. J. (2007) Identification of minor inner-membrane components of the Shigella type III secretion system 'needle complex'. Microbiology. 153: 2405–2415. https://doi.org/10.1099/mic.0.2007/007781-0 | |
| pKD46 | | Datsenko K.A., Wanner B.L. (2000) One-step inactivation of chromosomal genes in Escherichia coli K-12 using PCR products. Proc Natl Acad Sci U S A. 97: 6640-6645. https://doi.org/10.1073/pnas.120163297 | |
| pCP20 | | Datsenko K.A., Wanner B.L. (2000) One-step inactivation of chromosomal genes in Escherichia coli K-12 using PCR products. Proc Natl Acad Sci U S A. 97: 6640-6645. https://doi.org/10.1073/pnas.120163297 | |
| pKD3 | | Datsenko K.A., Wanner B.L. (2000) One-step inactivation of chromosomal genes in Escherichia coli K-12 using PCR products. Proc Natl Acad Sci U S A. 97: 6640-6645. https://doi.org/10.1073/pnas.120163297 | |
| pSB3292 | pBAD24, HilA | Lara-Tejero, M., Kato, J., Wagner, S., Liu, X., & Galán, J. E. (2011). A sorting platform determines the order of protein secretion in bacterial type III systems. Science (New York, NY), | |

| | | | |
|---|---|---|---|
| | | 331(6021), 1188–1191. http://doi.org/10.1126/science.1201476 | |
| pSUP-pBpa | | Ryu, Y., & Schultz, P. G. (2006). Efficient incorporation of unnatural amino acids into proteins in Escherichia coli. Nature Methods, 3(4), 263–265. http://doi.org/10.1038/nmeth864 | |
| pMIB6089 | pT10, SctRSTU$_{N258A}^{FLAG}$ | Dietsche, T., Mebrhatu, M. T., Brunner, M. J., Abrusci, P., Yan, J., Franz-Wachtel, M., ... & Kohlbacher, O. (2016). Structural and functional characterization of the bacterial type III secretion export apparatus. PLoS pathogens, 12(12), e1006071. http://doi:10.1371/journal.ppat.1006071 | |
| pMIB5272 | pT10, SctU$_{N258A}^{FLAG}$ | Monjarás Feria, J. V., Lefebre, M. D., Stierhof, Y.-D., Galán, J. E., & Wagner, S. (2015). Role of autocleavage in the function of a type III secretion specificity switch protein in *Salmonella enterica* serovar Typhimurium. *mBio*, *6*(5), e01459–15. http://doi.org/10.1128/mBio.01459-15 | |
| pSB3598 | pT10, SctU | Wagner, S., Königsmaier, L., Lara-Tejero, M., Lefebre, M., Marlovits, T. C., & Galán, J. E. (2010). Organization and coordinated assembly of the type III secretion export apparatus. Proceedings of the National Academy of Sciences of the United States of America, 107(41), 17745–17750. http://doi.org/10.1073/pnas.1008053107 | |
| pMIB7234 | pT10, SctRS$_{T38X}$TU$_{N258A}^{FLAG}$ | this study | Made by *in vivo* assembly of the PCR product of the following primer/template pair: Primers 46 and 47 from pMIB6089. |
| pMIB7235 | pT10, SctRS$_{V39X}$TU$_{N258A}^{FLAG}$ | this study | Made by *in vivo* assembly of the PCR product of the following primer/template pair: Primers 48 and 49 from pMIB6089. |
| pMIB7236 | pT10, SctRS$_{T40X}$TU$_{N258A}^{FLAG}$ | this study | Made by *in vivo* assembly of the PCR product of the following primer/template pair: Primers 50 and 51 from pMIB6089. |
| pMIB7237 | pT10, SctRS$_{Q41X}$TU$_{N258A}^{FLAG}$ | this study | Made by *in vivo* assembly of the PCR product of the following primer/template pair: Primers 52 and 53 from pMIB6089. |
| pMIB7027 | pT10, SctRS$_{L42X}$TU$_{N258A}^{FLAG}$ | this study ??? Rebecca | Made by *in vivo* assembly of the PCR product of the following primer/template pair: Primers 54 and 55 from pMIB6089. |
| pMIB6693 | pT10, SctRSTU$_{A94X, N258A}^{FLAG}$ | this study | Made by *in vivo* assembly of the PCR product of the following |

| | | | |
|---|---|---|---|
| | | | primer/template pair: Primers 56 and 57 from pMIB6089. |
| pMIB7201 | pT10, $SctRSTU_{T107X, N258A}^{FLAG}$ | this study | Made by *in vivo* assembly of the PCR product of the following primer/template pair: Primers 58 and 59 from pMIB6089. |
| pMIB7202 | pT10, $SctRSTU_{A109X, N258A}^{FLAG}$ | this study | Made by *in vivo* assembly of the PCR product of the following primer/template pair: Primers 60 and 61 from pMIB6089. |
| pMIB7203 | pT10, $SctRSTU_{L110X, N258A}^{FLAG}$ | this study | Made by *in vivo* assembly of the PCR product of the following primer/template pair: Primers 62 and 63 from pMIB6089. |
| pMIB7205 | pT10, $SctRSTU_{S115X, N258A}^{FLAG}$ | this study | Made by *in vivo* assembly of the PCR product of the following primer/template pair: Primers 64 and 65 from pMIB6089. |
| pMIB7206 | pT10, $SctRSTU_{N118X, N258A}^{FLAG}$ | this study | Made by *in vivo* assembly of the PCR product of the following primer/template pair: Primers 66 and 67 from pMIB6089. |
| pMIB7207 | pT10, $SctRSTU_{V120X, N258A}^{FLAG}$ | this study | Made by *in vivo* assembly of the PCR product of the following primer/template pair: Primers 68 and 69 from pMIB6089. |
| pMIB7204 | pT10, $SctRSTU_{F127X, N258A}^{FLAG}$ | this study | Made by *in vivo* assembly of the PCR product of the following primer/template pair: Primers 70 and 71 from pMIB6089. |
| pMIB6916 | pT10, $SctU_{A149X}$ | this study | Made by *in vivo* assembly of the PCR product of the following primer/template pair: Primers 86 and 87 from pSB3598. |
| pMIB6917 | pT10, $SctU_{I171X}$ | this study | Made by *in vivo* assembly of the PCR product of the following |

| | | | |
|---|---|---|---|
| | | | primer/template pair: Primers 88 and 89 from pSB3598. |
| pMIB7021 | pT10, $SctU_{A190X}$ | this study | Made by *in vivo* assembly of the PCR product of the following primer/template pair: Primers 94 and 95 from pSB3598. |
| pMIB6695 | pT10, $SctU_{W175X}$ | this study | Made by *in vivo* assembly of the PCR product of the following primer/template pair: Primers 90 and 91 from pSB3598. |
| pMIB6694 | pT10, $SctU_{L182X}$ | this study | Made by *in vivo* assembly of the PCR product of the following primer/template pair: Primers 92 and 93 from pSB3598. |
| pMIB6696 | pT10, $SctU_{A94X}$ | this study | Made by *in vivo* assembly of the PCR product of the following primer/template pair: Primers 56 and 57 from pSB3598. |
| pMIB7027 | pT10, $SctRS_{L42X}TU_{N258A}{}^{FLAG}$ | this study | Made by *in vivo* assembly of the PCR product of the following primer/template pair: Primers 54 and 55 from pMIB6089. |
| pMIB7026 | pT10, $SctRS_{L47X}TU_{N258A}{}^{FLAG}$ | this study | Made by *in vivo* assembly of the PCR product of the following primer/template pair: Primers 96 and 97 from pMIB6089. |
| pMIB7023 | pT10, $SctRS_{G50X}TU_{N258A}{}^{FLAG}$ | this study | Made by *in vivo* assembly of the PCR product of the following primer/template pair: Primers 98 and 99 from pMIB6089. |
| pMIB7022 | pT10, $SctRS_{L54X}TU_{N258A}{}^{FLAG}$ | this study | Made by *in vivo* assembly of the PCR product of the following primer/template pair: Primers 100 and 101 from pMIB6089. |
| pMIB7025 | pT10, $SctRS_{C57X}TU_{N258A}{}^{FLAG}$ | this study | Made by *in vivo* assembly of the PCR product of the following |

| | | | |
|---|---|---|---|
| | | | primer/template pair: Primers 102 and 103 from pMIB6089. |
| pMIB7032 | pT10, $SctRS_{F61X}TU_{N258A}^{FLAG}$ | this study | Made by *in vivo* assembly of the PCR product of the following primer/template pair: Primers 104 and 105 from pMIB6089. |
| pMIB7031 | pT10, $SctRS_{E69X}TU_{N258A}^{FLAG}$ | this study | Made by *in vivo* assembly of the PCR product of the following primer/template pair: Primers 106 and 107 from pMIB6089. |
| pMIB7030 | pT10, $SctRST_{S186X}U_{N258A}^{FLAG}$ | this study | Made by *in vivo* assembly of the PCR product of the following primer/template pair: Primers 108 and 109 from pMIB6089. |
| pMIB7029 | pT10, $SctRST_{L190X}U_{N258A}^{FLAG}$ | this study | Made by *in vivo* assembly of the PCR product of the following primer/template pair: Primers 110 and 111 from pMIB6089. |
| pMIB7028 | pT10, $SctRST_{F198X}U_{N258A}^{FLAG}$ | this study | Made by *in vivo* assembly of the PCR product of the following primer/template pair: Primers 112 and 113 from pMIB6089. |
| pMIB7033 | pT10, $SctRST_{L194X}U_{N258A}^{FLAG}$ | this study | Made by *in vivo* assembly of the PCR product of the following primer/template pair: Primers 114 and 115 from pMIB6089. |
| pMIB7024 | pT10, $SctR_{L211X}STU_{N258A}^{FLAG}$ | this study | Made by *in vivo* assembly of the PCR product of the following primer/template pair: Primers 116 and 117 from pMIB6089. |
| pMIB7287 | pT10, $SctRSTU_{F24X, N258A}^{FLAG}$ | this study | Made by *in vivo* assembly of the PCR product of the following primer/template pair: Primers 118 and 119 from pMIB6089. |
